# Enhancing Patient Lymphocyte Response to Peritoneal Malignancies Using a Personalized Immunocompetent Microfluidic Co-Culture Platform

**DOI:** 10.64898/2026.02.19.706377

**Authors:** Cecilia R. Schaaf, Damian C. Hutchins, Tiefu Liu, Mitra Kooshki, Calvin J. Wagner, Nicholas Edenhoffer, Nadeem Wajih, Steven D. Forsythe, Robyn Geissinger, Edward A. Levine, Perry Shen, Pierre Triozzi, Lance D. Miller, Adam R. Hall, Shay Soker, Konstantinos I. Votanopoulos

**Affiliations:** Department of Pathology, Wake Forest University School of Medicine, Winston-Salem, NC, USA; Wake Forest Organoid Research Center, Wake Forest University School of Medicine, Winston-Salem, NC, USA; Wake Forest Institute for Regenerative Medicine, Wake Forest University School of Medicine, Winston-Salem, NC, USA; Department of Biomedical Engineering, Wake Forest University School of Medicine, Winston-Salem, NC, USA; Department of Surgery, Section on Surgical Oncology, Wake Forest University School of Medicine, Winston-Salem, NC, USA; Department of Hematology and Oncology, Wake Forest University School of Medicine, Winston-Salem, NC, USA; Department of Cancer Biology, Wake Forest University School of Medicine, Winston-Salem, NC, USA

## Abstract

Harnessing patient immune cells via adoptive cellular therapy is a promising cancer therapeutic strategy. However, major challenges remain for advanced solid malignancies, including difficulty isolating sufficient tumor infiltrating lymphocytes (TILs) and limited targeting of diverse neoantigens in heterogenous tumors. To address this, we have developed a tumor-on-a-chip platform with co-cultured patient-derived tumor cells, autologous peripheral blood mononuclear cells (PBMCs), and lymphoid tissue-derived antigen presenting cells. This approach generates organoid interacting lymphocytes (OILs) with enhanced anti-tumor activity. In peritoneal malignancies, OIL-induced cytotoxicity of patient-matched tumor cells surpasses both TILs and static-expanded PBMCs. We find that this improved performance was linked to increased CD8^+^ T and NK cells among OILs, and increased effector cytokine polyfunctionality, particularly Granzyme A. Our platform represents a versatile and scalable approach to generate patient-specific therapeutic lymphocytes even when TILs are insufficient, offering a promising avenue to treat diverse solid tumors associated with poor outcomes under current immunotherapies.

**Teaser:** A tumor-on-a-chip device primes patient immune cells with tumor recognition for personalized immunotherapy applications.

## Introduction

Adoptive cell therapy (ACT) is an emergent form of immunotherapy treatment for cancer patients that involves isolating patient immune cells, expanding and potentially modifying their activity *ex vivo* to improve function and tumor targeting capability, and then transfusing the cells into the patient as a therapy. One particular ACT modality in which autologous tumor infiltrating lymphocytes (TILs) are isolated from surgically-resected or biopsied tumor tissue has demonstrated encouraging clinical response in several solid tumor types including melanoma (*1*), non-small cell lung cancer (*2*), and cervical cancer (*3*). While a positive response has been achieved for some individuals, consistent generation of a robust TIL population from all patients remains elusive. For many solid tumors, suppressive tumor microenvironment factors such as dense fibrosis, increased interstitial pressure, or suppressive regulatory immune cells can substantially reduce initial lymphocyte infiltration into the tumor and therefore negate TIL isolation efforts. Of the lymphocytes that are able to infiltrate the tumor, protracted exposure to the immune-inhibitory tumor microenvironment can lead to cellular exhaustion, thereby depleting TIL efficacy for cytokine production and cytolytic activity, even with *ex vivo* manipulation (*4, 5*). Finally, T cell receptors in isolated and expanded TILs only reflect tumor neoantigens that have been previously encountered, making them ineffectual for heterogenous tumors with diverse intra-tumor neoantigen populations (*6*).

As bio-fabrication capabilities have progressed, cancer research has increasingly turned toward patient-derived tumor culture as an *ex vivo* platform to recapitulate native tumor properties. For example, surgically-resected or biopsied cell isolates can be cultured within three-dimensional (3D) hydrogels that mimic the extracellular matrix (ECM). Such patient-derived models have emerged as a relatively low-cost, high-throughput platform for personalized preclinical testing of chemotherapy treatments (*7*), *ex vivo* evaluation of CAR-T cell therapy efficacy (*8*), and preclinical study of immunotherapy efficacy in appendiceal cancer (*9*), Merkel cell carcinoma (*10*), and melanoma (*11*).

Towards applying a patient tumor culture platform to address challenges encountered in current TIL therapy and extend ACT to a broader array of tumors, we developed a novel, biomimetic tumor-on-a-chip microfluidic device. Herein, autologous peripheral blood mononuclear cells (PBMCs) were circulated among tumor cells grown on ECM, enriched with patient-specific, secondary lymphoid tissue-derived presenting cells (APCs). This format promotes interaction between circulating PBMCs and ECM-adherent tumor/APC cells, resulting in a population of tumor-reactive T cells that we termed organoid interacting lymphocytes (OILs). In the present work, we demonstrate the potential of this device-based approach to prime therapeutic lymphocytes and improve ACT. We conduct *ex vivo* efficacy testing as a preliminary step to additional *in vivo* preclinical testing. As an initial model for this development, we employed clinical tissue specimens from patients with stage IV peritoneal cancers, associated with aggressive tumors that feature high clonal heterogeneity, dense fibrosis, and multiple site dissemination within the peritoneal cavity. These features not only make peritoneal malignancies more resistant to traditional chemo- and immunotherapy treatments (*12*)(*13*), but also make them particularly challenging targets for conventional TIL therapy. Our results establish OIL phenotype, reactivity, and *ex vivo* cytotoxicity against patient-matched tumors, thereby validating our microfluidic tumor-on-a-chip platform for on-demand generation of tumor-specific, therapeutic OILs. These investigations serve as the initial validation of a novel, tumor-agnostic ACT modality with potential applicability to a broad array of cancers.

## Results

### Patient Characteristics

Tumor tissue was collected from multiple spatially-distinct and heterogeneous abdominal lesions in individual patients undergoing cytoreductive surgery combined with hyperthermic intraperitoneal chemotherapy (HIPEC). Tumor specimens for our experiments were collected prior to the start of HIPEC. From each of these patients, matched secondary lymphoid tissue (lymph node or spleen) and peripheral blood were also collected. The total cohort included patients treated for either appendiceal cancer with dissemination in the peritoneal cavity (n=21) or primary peritoneal mesothelioma (n=10). Pertinent patient characteristics including pathologic diagnosis and prior treatments are included in Table 1. Female patients represented 61% (19/31) of the entire cohort. Due to the rarity of both malignancies and to minimize risk of patient identification, individual patient gender and age have been omitted.

**Table 1.**
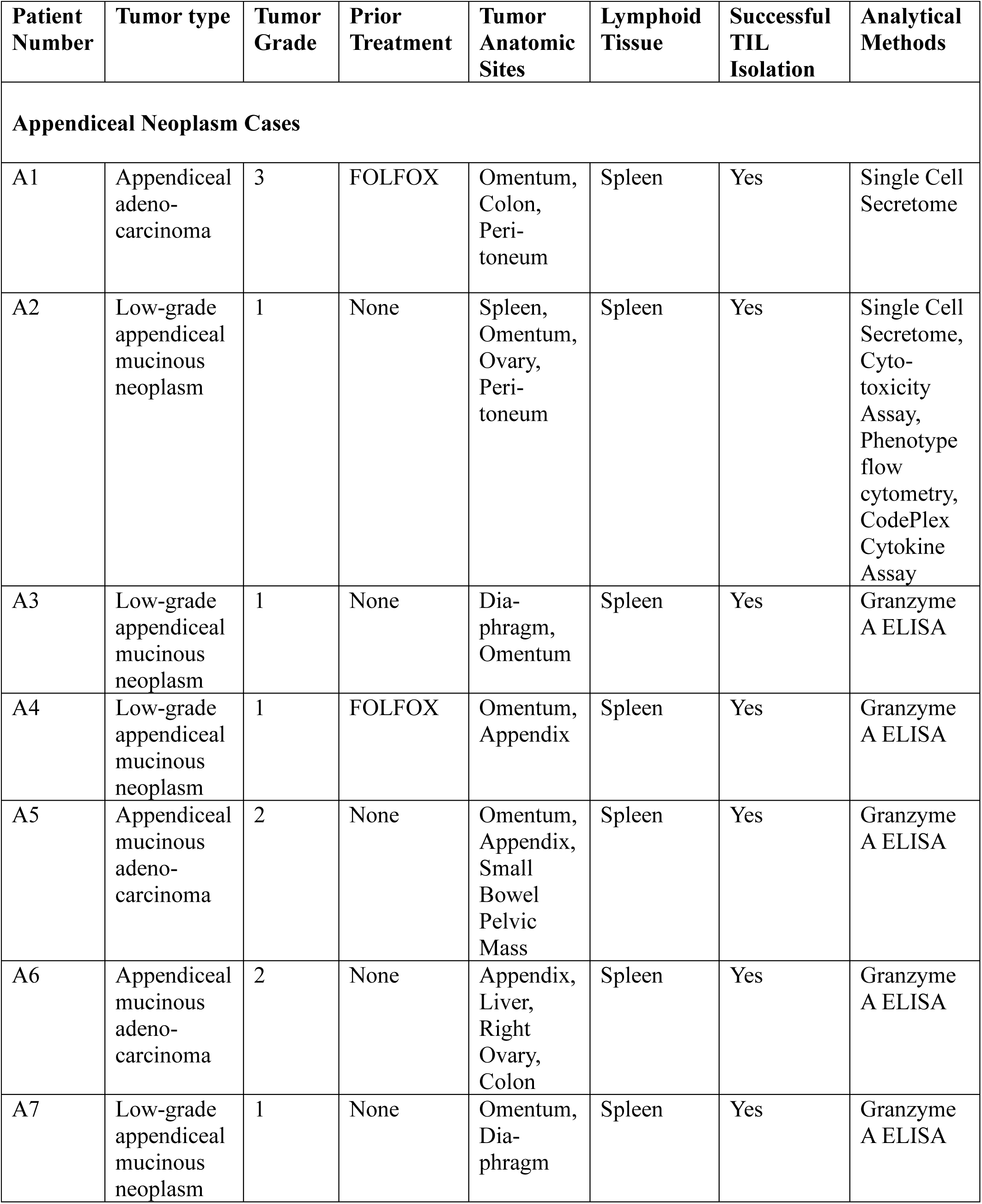
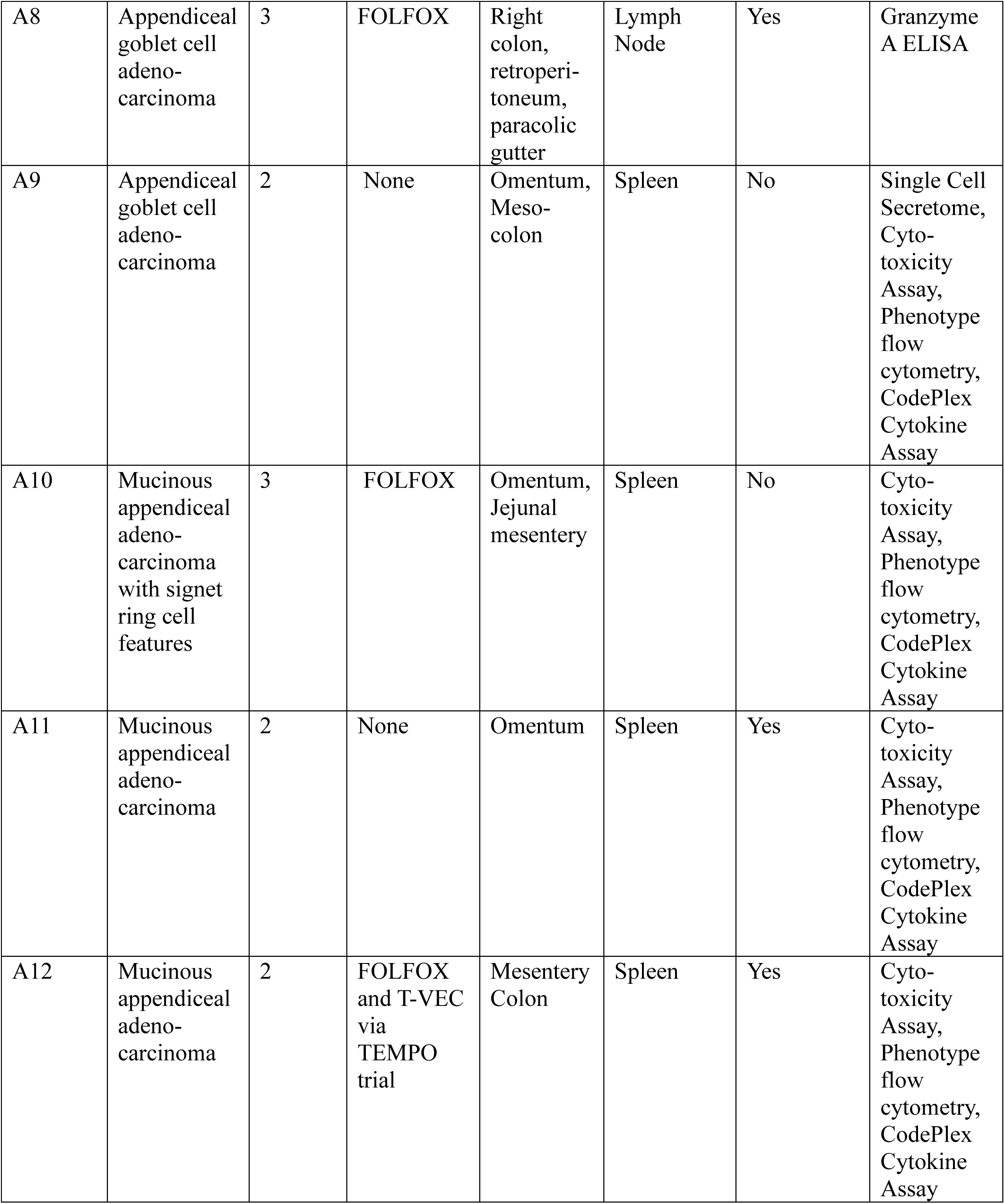
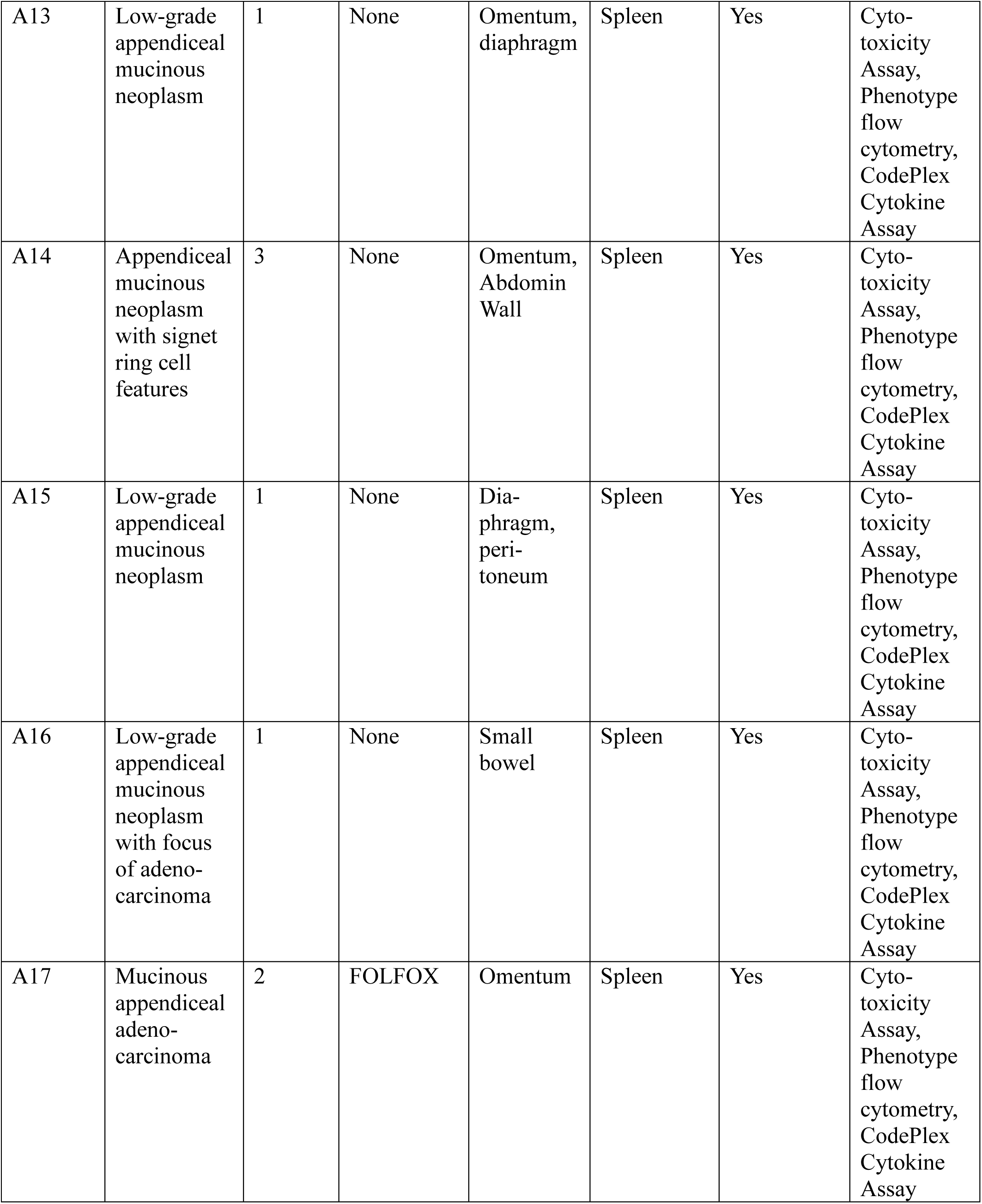
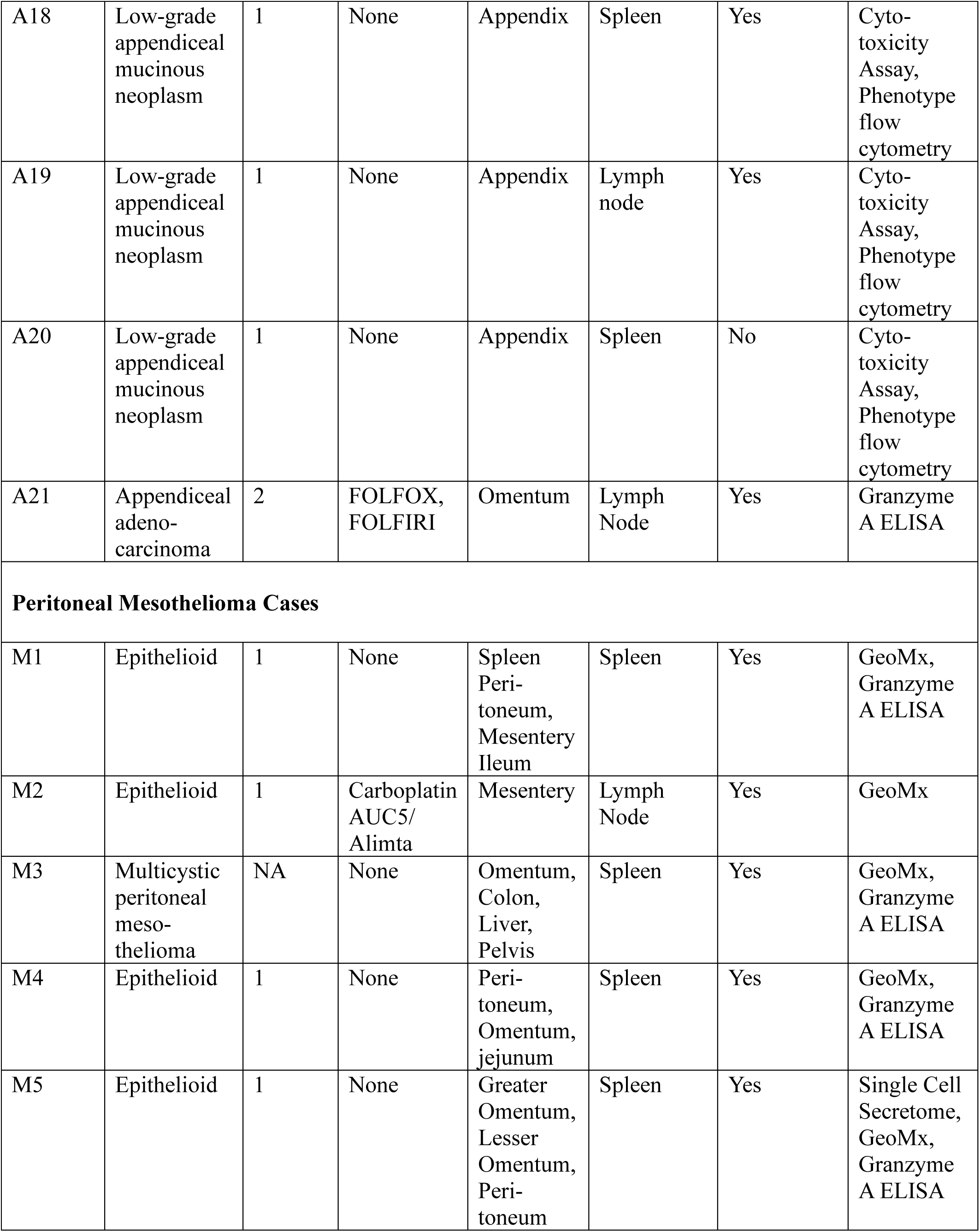
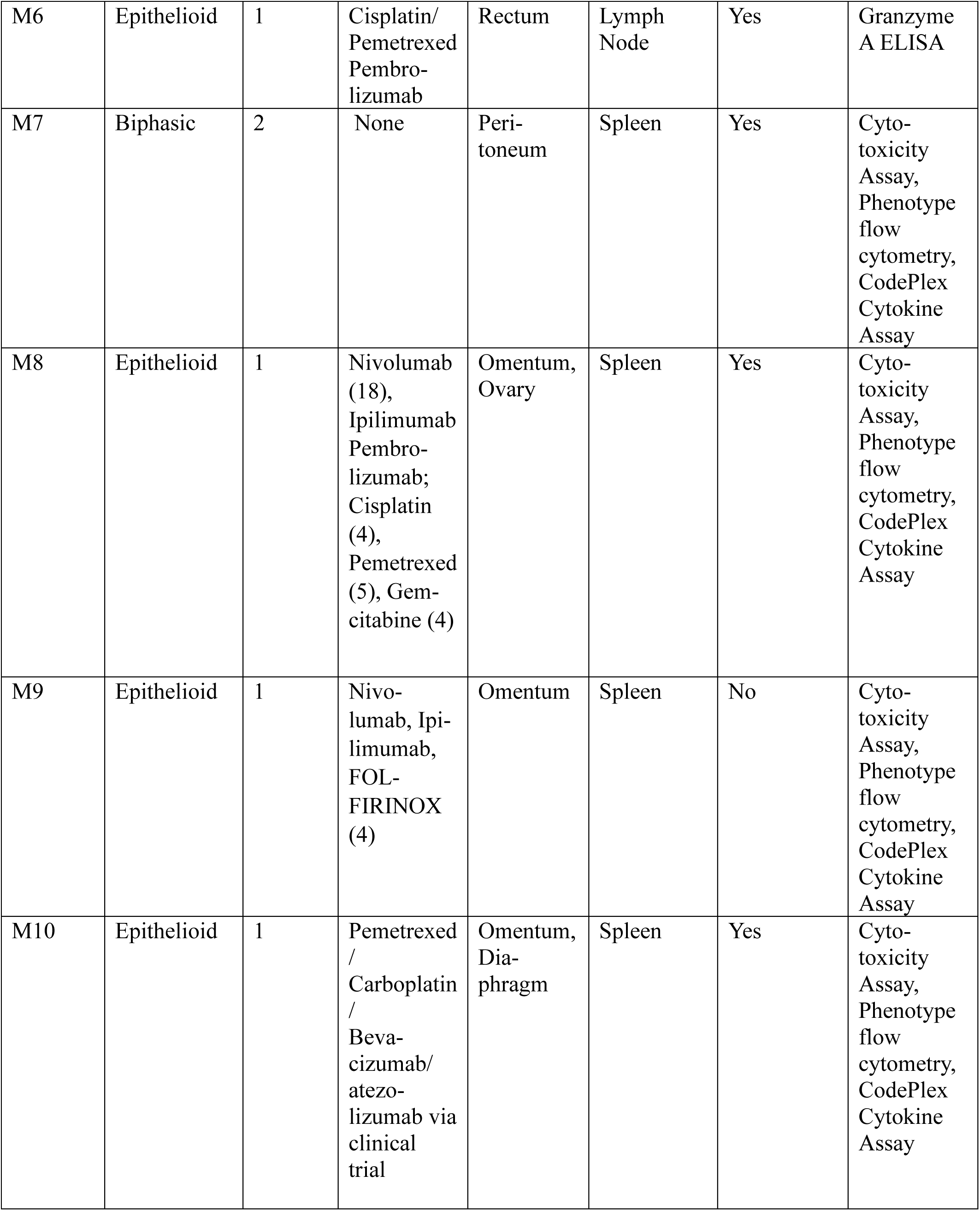
Patient Characteristics. Patient table describing tumor type and grade, previous treatment history, tumor resection anatomic sites, the source of secondary lymphoid tissue used for co-culture with tumor (normal spleen or normal lymph node, if spleen is indicated then the patient was already undergoing splenectomy due to tumor burden), whether tumor infiltrating lymphocytes (TILs) could be isolated from the resected tumors, and the analytical methods applied on the generated OILs, PBMCes, and TILs.

| Patient Number | Tumor type | Tumor Grade | Prior Treatment | Tumor Anatomic Sites | Lymphoid Tissue | Successful TIL Isolation | Analytical Methods |
| --- | --- | --- | --- | --- | --- | --- | --- |
| **Appendiceal Neoplasm Cases** |  |  |  |  |  |  |  |
| A1 | Appendiceal adeno-carcinoma | 3 | FOLFOX | Omentum, Colon, Peri-toneum | Spleen | Yes | Single Cell Secretome |
| A2 | Low-grade appendiceal mucinous neoplasm | 1 | None | Spleen, Omentum, Ovary, Peri-toneum | Spleen | Yes | Single Cell Secretome, Cytotoxicity Assay, Phenotype flow cytometry, CodePlex Cytokine Assay |
| A3 | Low-grade appendiceal mucinous neoplasm | 1 | None | Dia-phragm, Omentum | Spleen | Yes | Granzyme A ELISA |
| A4 | Low-grade appendiceal mucinous neoplasm | 1 | FOLFOX | Omentum, Appendix | Spleen | Yes | Granzyme A ELISA |
| A5 | Appendiceal mucinous adeno-carcinoma | 2 | None | Omentum, Appendix, Small Bowel Pelvic Mass | Spleen | Yes | Granzyme A ELISA |
| A6 | Appendiceal mucinous adeno-carcinoma | 2 | None | Appendix, Liver, Right Ovary, Colon | Spleen | Yes | Granzyme A ELISA |
| A7 | Low-grade appendiceal mucinous neoplasm | 1 | None | Omentum, Dia-phragm | Spleen | Yes | Granzyme A ELISA |
| A8 | Appendiceal goblet cell adeno-carcinoma | 3 | FOLFOX | Right colon, retroperitoneum, paracolic gutter | Lymph Node | Yes | Granzyme A ELISA |
| A9 | Appendiceal goblet cell adeno-carcinoma | 2 | None | Omentum, Meso-colon | Spleen | No | Single Cell Secretome, Cytotoxicity Assay, Phenotype flow cytometry, CodePlex Cytokine Assay |
| A10 | Mucinous appendiceal adeno-carcinoma with signet ring cell features | 3 | FOLFOX | Omentum, Jejunal mesentery | Spleen | No | Cytotoxicity Assay, Phenotype flow cytometry, CodePlex Cytokine Assay |
| A11 | Mucinous appendiceal adeno-carcinoma | 2 | None | Omentum | Spleen | Yes | Cytotoxicity Assay, Phenotype flow cytometry, CodePlex Cytokine Assay |
| A12 | Mucinous appendiceal adeno-carcinoma | 2 | FOLFOX and T-VEC via TEMPO trial | Mesentery Colon | Spleen | Yes | Cytotoxicity Assay, Phenotype flow cytometry, CodePlex Cytokine Assay |
| A13 | Low-grade appendiceal mucinous neoplasm | 1 | None | Omentum, diaphragm | Spleen | Yes | Cyto-toxicity Assay, Phenotype flow cytometry, CodePlex Cytokine Assay |
| A14 | Appendiceal mucinous neoplasm with signet ring cell features | 3 | None | Omentum, Abdomin Wall | Spleen | Yes | Cyto-toxicity Assay, Phenotype flow cytometry, CodePlex Cytokine Assay |
| A15 | Low-grade appendiceal mucinous neoplasm | 1 | None | Dia-phragm, peri-toneum | Spleen | Yes | Cyto-toxicity Assay, Phenotype flow cytometry, CodePlex Cytokine Assay |
| A16 | Low-grade appendiceal mucinous neoplasm with focus of adeno-carcinoma | 1 | None | Small bowel | Spleen | Yes | Cyto-toxicity Assay, Phenotype flow cytometry, CodePlex Cytokine Assay |
| A17 | Mucinous appendiceal adeno-carcinoma | 2 | FOLFOX | Omentum | Spleen | Yes | Cyto-toxicity Assay, Phenotype flow cytometry, CodePlex Cytokine Assay |
| A18 | Low-grade appendiceal mucinous neoplasm | 1 | None | Appendix | Spleen | Yes | Cyto-toxicity Assay, Phenotype flow cytometry |
| A19 | Low-grade appendiceal mucinous neoplasm | 1 | None | Appendix | Lymph node | Yes | Cyto-toxicity Assay, Phenotype flow cytometry |
| A20 | Low-grade appendiceal mucinous neoplasm | 1 | None | Appendix | Spleen | No | Cyto-toxicity Assay, Phenotype flow cytometry |
| A21 | Appendiceal adenocarcinoma | 2 | FOLFOX, FOLFIRI | Omentum | Lymph Node | Yes | Granzyme A ELISA |
| Peritoneal Mesothelioma Cases |  |  |  |  |  |  |  |
| M1 | Epithelioid | 1 | None | Spleen<br>Peri-toneum,<br>Mesentery<br>Ileum | Spleen | Yes | GeoMx,<br>Granzyme<br>A ELISA |
| M2 | Epithelioid | 1 | Carboplatin<br>AUC5/<br>Alimta | Mesentery | Lymph<br>Node | Yes | GeoMx |
| M3 | Multicystic peritoneal mesothelioma | NA | None | Omentum,<br>Colon,<br>Liver,<br>Pelvis | Spleen | Yes | GeoMx,<br>Granzyme<br>A ELISA |
| M4 | Epithelioid | 1 | None | Peri-toneum,<br>Omentum,<br>jejunum | Spleen | Yes | GeoMx,<br>Granzyme<br>A ELISA |
| M5 | Epithelioid | 1 | None | Greater<br>Omentum,<br>Lesser<br>Omentum,<br>Peri-toneum | Spleen | Yes | Single Cell<br>Secretome,<br>GeoMx,<br>Granzyme<br>A ELISA |
| M6 | Epithelioid | 1 | Cisplatin/<br>Pemetrexed<br>Pembro-<br>lizumab | Rectum | Lymph<br>Node | Yes | Granzyme<br>A ELISA |
| M7 | Biphasic | 2 | None | Peri-toneum | Spleen | Yes | Cyto-<br>toxicity<br>Assay,<br>Phenotype<br>flow<br>cytometry,<br>CodePlex<br>Cytokine<br>Assay |
| M8 | Epithelioid | 1 | Nivolumab<br>(18),<br>Ipilimumab<br>Pembro-<br>lizumab;<br>Cisplatin<br>(4),<br>Pemetrexed<br>(5), Gem-<br>citabine (4) | Omentum,<br>Ovary | Spleen | Yes | Cyto-<br>toxicity<br>Assay,<br>Phenotype<br>flow<br>cytometry,<br>CodePlex<br>Cytokine<br>Assay |
| M9 | Epithelioid | 1 | Nivo-<br>lumab, Ipi-<br>limumab,<br>FOL-FIRINOX<br>(4) | Omentum | Spleen | No | Cyto-<br>toxicity<br>Assay,<br>Phenotype<br>flow<br>cytometry,<br>CodePlex<br>Cytokine<br>Assay |
| M10 | Epithelioid | 1 | Pemetrexed<br>/<br>Carboplatin<br>/<br>Beva-<br>cizumab/<br>atezo-<br>lizumab via<br>clinical<br>trial | Omentum,<br>Dia-phragm | Spleen | Yes | Cyto-<br>toxicity<br>Assay,<br>Phenotype<br>flow<br>cytometry,<br>CodePlex<br>Cytokine<br>Assay |

### Microfluidic Chip Bio-fabrication and Experimental Timeline

Cells derived from patient tumor and APC-containing lymphoid tissue were co-cultured in hydrogel within the central chamber of a microfluidic chip (Figure 1A -C; also see *Materials and Methods*). A reservoir of autologous PBMCs, suspended in activation/expansion media, was then connected to the chip inlet (Figure 1A) and circulated continuously through the tumor-APC co-culture for 7 days via peristaltic pumping to yield OILs (Figure 1A and D). To evaluate the effect of circulation, a portion of each patient PBMCs were separately cultured in static conditions with activation/expansion media and in the absence of tumor cells to generate expanded PBMCs (PBMCes; Figure 1D). As an additional control, tumor tissues were also cultured alone in static conditions to isolate naturally-occurring TILs (Figure 1D). As an FDA-approved ACT for melanoma, TILs served to provide a clinically-relevant comparison to OIL activity. Seven days after tissue collection, all three cell types (OILs, PBMCes, and TILs) were further expanded in static culture for an additional 7 days to increase cell numbers and then assessed for phenotype, functionality, and induction of tumor cell cytotoxicity *ex vivo* (Figure 1D). The total time from sample collection/cell isolation to final tumor cytotoxicity testing was 16 days.

**Figure 1.**
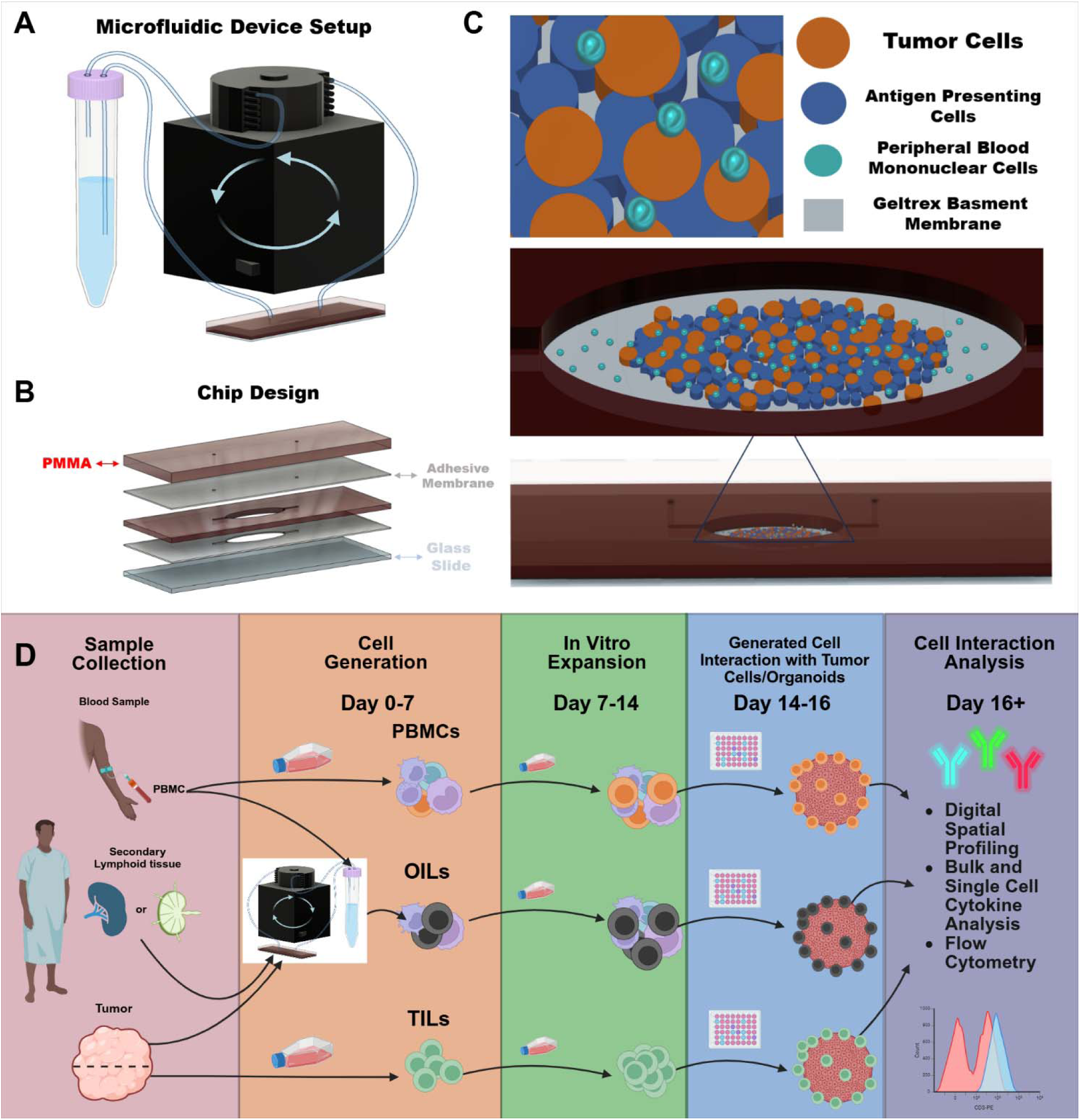
Experimental design and microfluidic chip schema. **(A)** Representation of the 75x26x6mm microfluidic chip model with inlet and outlet tubing connected to an MP2 Meinhardt peristaltic pump continuously circulating PBMCs through the chip from the 15mL conical reservoir. **(B)** Microfluidic chip device was assembled from 2 polymethyl methacrylate (PMMA) layers, 2 adhesive membrane layers, and a glass slide to form a central chamber with inlet and outlet ports on either end. **(C)** 3D modeled images depict where tumor cells (orange), secondary lymphoid tissue cells [from spleen or lymph node, containing antigen presenting cells (APCs), navy], and PBMCs (cyan) are located within the microfluidic chip central chamber. Tumor cells and secondary lymphoid tissue cells were seeded directly onto basement membrane (gray) prior to initiation of PBMC circulation. **(D)** While undergoing cytoreductive surgery, a portion of the tumor, healthy secondary lymphoid tissue and whole blood were collected from each patient. In the initial cell generation phase (Day 0-7), patient-matched peripheral blood mononuclear cells (PBMCs) were isolated from whole blood and cultured in either static culture conditions to create expanded PBMCs (PBMCes) or circulated through the microfluidic chip device, containing resected tumor and lymphoid tissue cells, to produce organoid interacting lymphocytes (OILs). Separately, naturally occurring tumor infiltrating lymphocytes (TILs) were isolated from a portion of the same resected tumor tissue. Subsequently, all groups were further expanded in static culture conditions to increase cell numbers (Days 7-14). The different immune cell groups were then independently co-cultured with patient-matched tumor cells seeded on basement membrane or grown as tumor organoids (Day 14) to determine relative tumor cytotoxicity and immune cell effector propensity. Figure created using Biorender software.

### OILs Demonstrate Increased Tumor Cell Cytotoxicity over TILS and expanded PBMCs

We first compared the efficacy of immune cell-mediated tumor cell death (TCD) between OIL, PBMCe, and TIL treatments by co-culturing each population with patient-matched tumor cells in 2D culture for 24 hours. As a control for spontaneous cell death, tumor cells were also cultured alone. Phase contrast images qualitatively confirmed the relative interactions of each immune cell population with tumor cells. As shown in the representative images of tumor and immune co-cultures (Figure 2A, patient designation A18), increased clustering of small immune cells surrounding large, rounded tumor cells was observed frequently in the OIL-treated co-cultures (red circles), along with a reduced count of adherent tumor cells. This suggested increased interactions between immune and tumor cells that preceded tumor cell detachment and eventual TCD. In contrast, co-culture of tumor with PBMCe or TILs featured clusters of adherent, living tumor cells (black arrows), despite the presence of numerous lymphocytes in their vicinity. This suggested a lack of tumor recognition in these populations.

**Figure 2.**
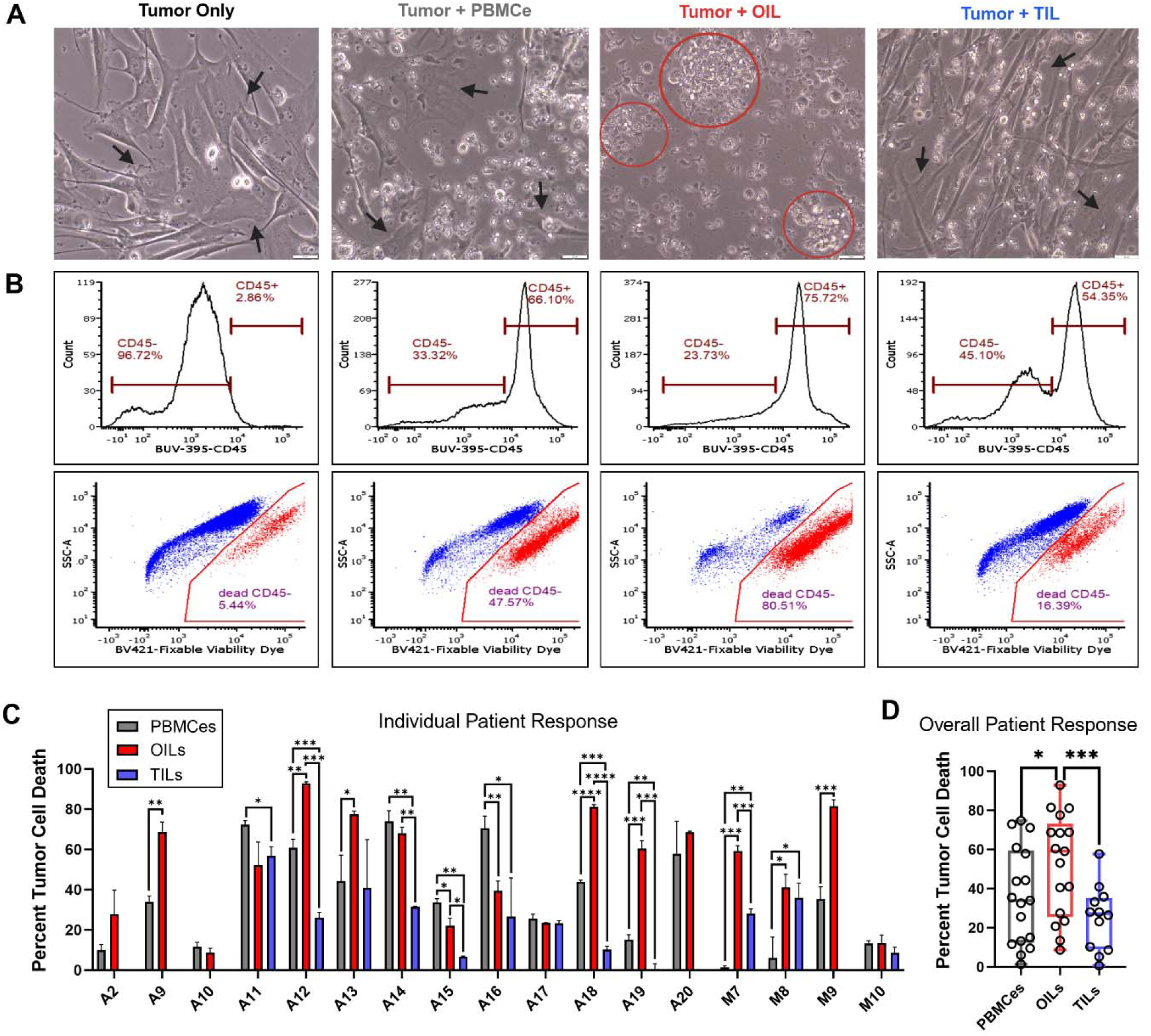
Treatment of peritoneal tumors with OILs resulted in increased tumor cell death over TILs and expanded PBMCs. **(A)** Representative images from tumor and immune cell [(PBMCe), (OIL), and (TIL)] co-cultures after 24-hour incubation to assess tumor cell death (TCD), sample from appendiceal patient A18. Black arrows indicate tumor cells, red circles indicate immune cell clusters. Scale bar 20µm (white). **(B)** Representative flow cytometry gating strategy from patient in (A) to quantify tumor cell (CD45^-^ cells in histogram) death via viability dye expression (scatterplot). **(C)** Individual patient response and **(D)** overall patient response measured by the percent of tumor cell death in co-culture with either PBMCes (gray), OILs (red), or TILs (blue). The percent of treatment-induced TCD was normalized to spontaneous cell death measured in the tumor cell only control cultures. Error bars represent standard deviation for each individual patient treatment group, measured by triplicate culture replicates. Boxplots represent the overall patient response (n=17 patients; 13 appendiceal and 4 mesothelioma, triplicate culture replicates for each patient, data points represent each patient mean response). Data in (D) were compared using linear mixed models and post hoc Tukey method while individual patient responses in (C) were compared using student T-test. *p< 0.05 **p< 0.01 ***p< 0.001 ****p<0.0001.

Co-culture treatment groups were further evaluated by quantitative flow cytometry analysis of their tumor cells (CD45^-^) (Figure 2B). For each immune cell population, the percent of TCD in its co-culture was determined and normalized to the spontaneous TCD observed in the matched tumor-only control culture (Figures 2C). Figure 2B shows the results of a gating strategy for first identifying CD45^-^ cells and then quantifying their viability based on live/dead staining. These results correspond to the same representative patient sample co-culture displayed in Figure 2A. For this patient (A18), the co-culture percentages of TCD were, for the tumor-only control and in the co-cultures, 47.6% for PBMCe, 80.5% for OIL, and 16.4% for TIL, while the TCD for the tumor-only control was only 5.4% (Figure 2B). Parallel images and flow cytometry gating strategy demonstrating similar increased activity of OILs in a representative mesothelioma patient (M8) are shown in Supplementary Figure S1A-B.

When TCD was determined for all patients combined (Figure 2D, overall patient analysis), OIL_TCD_ was significantly higher compared to both PBMCe and TIL (mean PBMCe 35.9%, OIL 52.2%, and TIL 24.9%; OILs vs PBMCe p<0.05 and OILs vs TILs p<0.001). When TCD was determined for all peritoneal malignancy patients individually (Figure 2C), we found that 47.1% (8/17) exhibited significantly higher OIL_TCD_ compared to corresponding PBMCe_TCD_ (Table 2, indicated by ^#^). In 29.4% (5/17) of patients, a sufficient number of TILs could not be derived for testing, further highlighting a major limitation of TILs as a therapeutic modality. In 64.71% (11/17) of patients, sufficient TIL numbers could be both isolated and expanded. However, in all comparisons of these patients, OIL_TCD_ was comparable or superior to that of TILs, reaching significance in 54.5% (6/11) of cases (Table 2, indicated by **^†^**).

**Table 2.** Mean tumor cell death values and comparisons for individual patient responses.

| Patient | PBMCE<br>mean TCD | OIL<br>mean TCD | TIL<br>mean TCD | OIL vs<br>PBMCE | OIL vs<br>TIL | PBMCE<br>vs TIL |
| --- | --- | --- | --- | --- | --- | --- |
| A2 | 9.55 | 27.42 | *insufficient cells* | NS | NC | NC |
| A9 | 33.98 | 68.68 | *insufficient cells* | # p<0.01 | NC | NC |
| A10 | 11.60 | 8.74 | *insufficient cells* | NS | NC | NC |
| A11 | 72.95 | 53.16 | 57.70 | NS | NS | p<0.05 |
| A12 | 60.96 | 92.90 | 26.45 | # p<0.01 | † p<0.001 | p<0.001 |
| A13 | 44.33 | 77.53 | 40.98 | # p<0.05 | NS | NC |
| A14 | 74.72 | 68.82 | 33.19 | NS | † p<0.001 | p<0.01 |
| A15 | 32.51 | 20.82 | 5.17 | p<0.001 | † p<0.05 | p<0.01 |
| A16 | 71.11 | 40.57 | 28.06 | p<0.01 | NS | p<0.05 |
| A17 | 25.60 | 23.48 | 23.24 | NS | NS | NS |
| A18 | 43.75 | 81.11 | 10.30 | # p<0.0001 | † p<0.0001 | p<0.001 |
| A19 | 15.05 | 60.44 | 0.6 | # p<0.001 | † p<0.001 | p<0.01 |
| A20 | 57.91 | 68.45 | *insufficient cells* | NS | NC | NC |
| M7 | 1.34 | 59.09 | 28.04 | # p<0.001 | † p<0.001 | p<0.01 |
| M8 | 6.01 | 41.05 | 35.88 | # p<0.05 | NS | p<0.05 |
| M9 | 35.48 | 81.47 | *insufficient cells* | # p<0.001 | NC | NC |
| M10 | 13.19 | 13.43 | 8.66 | NS | NS | NS |

Interestingly, PBMCe co-cultures induced significantly higher TCD than TILs in 58.3% (7/12) of relevant patients (Table 2) and even reached significantly higher than OILs in 2 of the 17 total patients (Table 2). This behavior could be attributed to prior *in vivo* interactions of the circulating peripheral immune cells with tumor neoantigens, potentially amplified by selection during the *ex vivo* expansion process. This raised the intriguing possibility that PBMC expansion alone could provide an alternative to TILs as a source of immune cells for ACT, especially in patients for whom TILs are unavailable due to low abundance, low activity, or an unresectable tumor. Still, the unexpected efficacy of PBMCes does not negate the value of OILs. Indeed, as already described above, comparing TCD between immune cell groups for all patients combined (Figure 2D), we found that OIL_TCD_ was significantly higher than either PBMCe_TCD_ or TIL_TCD_. The general trend was still observed even when considering the two primary tumor sources separately (appendiceal or mesothelioma, Supplementary Fig. S1C). Despite variation within each group stemming from patient heterogeneity, this enhanced TCD demonstrated a strong impact linked to the circulation process and a subsequent benefit in effectiveness.

### Microfluidic Chip Training Enriches for and Activates Effector Lymphocyte Populations

We next sought to determine the predominant immune phenotypes present in each cellular population and correlate the impact of phenotypes to observed TCD. Sixteen peritoneal malignancy cases were analyzed using flow cytometry to determine the phenotypic compositions of PBMCes, OILs, and TILs, and compared with matched baseline PBMCs (PBMCb) collected at initial PBMC isolation and analyzed in the same fashion (Figure 3A). Among all analyzed CD45^+^ cells, major immune cell phenotypes were identified as CD3^+^CD8^+^ T cells, CD3^+^CD4^+^ T cells, CD56^+^CD3^-^ natural killer (NK) cells, CD56^+^CD3^+^ NK T cells, CD20^+^ B cells, and “other” cells within CD45^+^ or CD3^+^ (including CD4^-^CD8^-^ and CD4^+^CD8^+^ T cells, TCRγδ T cells, monocytes, and dendritic cell populations), using the gating strategy described in Supplementary Figure S2.

**Figure 3.**
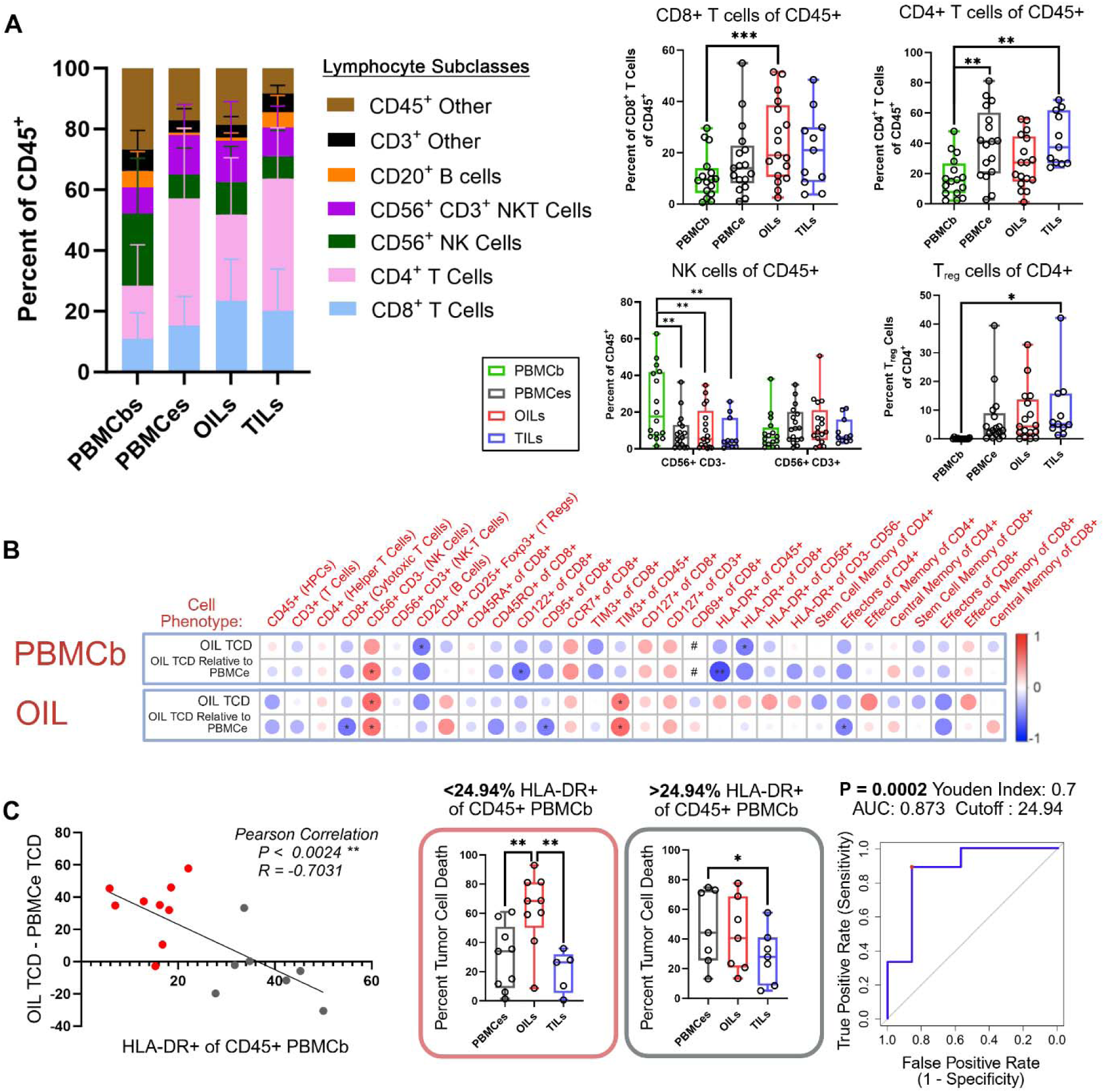
Immune cell population phenotypes can predict anti-tumor cytotoxic efficacy. **(A)** Bar graphs and box plots indicating the percent of predominant immune cell phenotypes present among groups [baseline PBMC (PBMCb), (PBMCe), (OIL), and (TIL)], determined by flow cytometry analysis. **(B)** Correlation heatmap depicting the normalized percent of tumor cell death (TCD) for both OILs and the difference between OIL TCD and PBMCe TCD (row titles in black, values derived from figure 2) compared to immune cell phenotype marker expression in PBMCbs and OILs (red column titles indicate phenotype measured via flow cytometry, red row titles indicate if these phenotypes are measured in PBMCbs or OILs). Measurements are compared by Pearson correlation coefficient. Heatmap color scale bar and size of the circle indicate the Pearson correlation coefficient for each relationship, with dark red large circles equal to 1 and dark blue large circles equal to -1. Significant relationships are indicated by *s, the # in PBMCb phenotype measurement indicates value not measured. **(C)** Pearson correlation plot comparing the percent of HLA-DR^+^CD45^+^ cells in the PBMCb population to the difference between OIL and PBMCe TCD, separated by low (red) versus high (gray) percent HLA-DR^+^ CD45^+^ PBMCbs (cutoff value of 24.9494%). Bar graphs show the percent TCD for PBMCes, OILs, and TILs for patients with HLA-DR+ of CD45+ PBMCbs below (red box) and above (gray box) the cutoff value of 24.9494%. Receiver-operator curve (ROC) demonstrating sensitivity and specificity of OIL – PBMCe TCD based on a cutoff value of 24.9494% for HLA-DR+ of CD45+ PBMCbs. Data points in box plots represent individual patients (n=16 patients). In all, box plots show minimum, median, and maximum values. Statistical significance calculated for linear mixed models using post hoc Tukey method. Pearson correlation coefficients analyzed using T-test. ROC curve calculated using Wilson/Brown method. *p<0.05 **p<0.01 ***p<0.001 ****p<0 .0001.

Upon first characterizing the general phenotypic responses within our immune cell populations, OILs showed a significant increase of CD8^+^ T cells compared to PBMCb (mean OILs 24.4% vs PBMCb 11.00% p<0.001; Figure 3A.). With respect to CD4^+^ T cells, PBMCes and TILs –– but not OILs –– showed a significant increase compared to PBMCb (mean PBMCb 17.4% vs PBMCe 40.0% p<0.01; and vs OILs 27.5% NS). Within the CD4^+^ T cell population, TILs showed a significant increase in T regulatory cell numbers (FOXP3^+^CD25^+^) over PBMCb (mean PBMCb 0.1% vs TILs 9.9% p<0.05). Regarding NK cells, PBMCes, OILs, and TILs all showed a significant decrease relative to the PBMCb population (mean PBMCb 24.1% vs PBMCe 88.9% p<0.01; vs OILs 10.6% p<0.01; and vs TILs 7.77% p<0.01; Figure 3A). Similar general trends were observed irrespective of the two primary tumor cohorts type (appendiceal or mesothelioma, Supplementary Fig. S3A-B). Collectively, these findings indicated that while there was an overall enrichment of CD8^+^ lymphocytes in OILs, there was also a patient-to-patient heterogeneity observed in the immune cell response when simply examining the proportions of cell phenotypes present. Consequently, further exploration of immune cell functions corresponding to TCD was warranted.

To address this, we, we initially assessed human leukocyte antigen (HLA-DR) expression as an indicator of antigen presentation and subsequent immune cell activation (Supplementary Fig. S4). We focused first on key cell populations with relevance to tumor cell cytotoxicity including: CD45^+^ cells, CD8^+^ T cells, and CD56^+^ NK cells – the latter two of which have been shown to have increased activation and enhanced tumor cytotoxic response when expressing HLA-DR (*14*),(*15*) – as well as the APC population (CD3^-^CD56^-^CD45^+^). In all cell populations, the OILs showed significantly higher expression compared to either PBMCb or TILs, but comparable to PBMCe (Supplementary Fig. S4). While it is possible the observed effects in OILs may be attributed to the activation and expansion media used during culturing, the fact that only OILs showed a significant increase in the abundance of CD8^+^ T cells (Figure 3A, top row) indicated that OILs likely contained a larger pool of activated effector cell types than other immune cell populations across all patients, shown previously to be important to immune-mediated tumor cytotoxicity (*16, 17*).

### Cytotoxic Efficacy of OILs can be Predicted by Baseline Phenotype Expression

To closer investigate the causes of patient-to-patient variability in OIL activation and efficacy (see Figure 2C) closer, we next sought to correlate the abundance of diverse immune cell phenotype markers in both PBMCb (Figure 3B, top) and OILs (Figure 3B, bottom) with OIL_TCD_ and PBMCe_TCD_ in each matched patient tumor. This was considered in two ways: OIL_TCD_ (upper rows) and as the relative difference in the TCDs between OILs and PBMCe (OIL_TCD_ - PBMCe_TCD_, lower rows) to capture the performance improvement realized by our tumor-on-a-chip training methodology relative to static expansion alone. These comparisons enabled us to assess if a particular cell phenotype could forecast differences in TCD, thus helping to predict which patients could benefit the most from OILs. Examining PBMCb phenotype correlation to OIL_TCD_ relative to PBMCe_TCD_, we found a significant negative correlation to the percentage of HLA-DR^+^ CD45^+^ T cells (-0.58, p<0.05), signifying that patients with a lower percentage of HLA-DR^+^ cells among their CD45^+^ PBMCs at the time of collection were likely to later exhibit high OIL_TCD_. Conversely, higher HLA-DR expression in CD45^+^ PBMCbs would lead to a static expanded PBMCe population that performed equal to matched OILs. This observation may further suggest that *in vitro* static expansion of PBMCb cells to produce PBMCes could specifically enrich HLA-DR^+^ CD45^+^ cells as an activated, tumor-specific effector cell population, thereby causing PBMCe_TCD_ to be high.

To better define the impact of peripheral blood HLA-DR^+^CD45^+^ cells on the efficacy of OILs compared to PBMCes, we next plotted (OIL_TCD_ - PBMCe_TCD_) against the abundance of HLA-DR^+^CD45^+^ cells among the PBMCb population (Figure 3C). From this, we found that the two variables were indeed highly correlated (*p*=0.0024), yielding a Pearson’s correlation of -0.7. This analysis also enabled us to classify patients based on their baseline HLA-DR abundance to predict the likely benefit of OILs production relative to simple static PBMC expansion in terms of resulting TCD. From this, we identified that patients with <24.94% HLA-DR^+^CD45^+^ PBMCb cells had significantly increased OIL_TCD_ compared to PBMCe_TCD_ (PBMCe 29.66% vs OILs 62.44% p<0.01; Figure 3C, red box) while those with higher amounts showed no difference between OIL_TCD_ and PBMCe_TCD_ (PBMCe 47.88% vs OILs 42.55% NS; Figure 3C, grey box). Evaluating this correlation via a receiver-operator curve (ROC), we found a sensitivity of 88.89% and a specificity of 85.71% (area under the curve = 0.873, p<0.0002; Figure 3C), demonstrating its predictive power.

Additional phenotypic analyses of the OILs population showed that the percentage of NK cells among OILs was positively correlated to both increased OIL_TCD_ (0.5959, p<0.505; Figure 3B, row 3) and to increased OIL_TCD_ relative to PBMCe_TCD_ (OIL_TCD_ - PBMCe_TCD_) (0.5959, p<0.505; Figure 3B, row 4). We also found a significant negative correlation between the percentage of CD122^+^CD8^+^ T cells and OIL_TCD_ - PBMCe_TCD_ (-0.59, p<0.05). In recent literature, this cell subset has been identified to act as either an indicator of an active anti-tumor response or a marker of immune suppression, dependent on context (*18*). Increased expression of T cell immunoglobulin and mucin domain 3 (TIM-3) on CD45^+^ OILs was also positively correlated to OIL_TCD_ (0.5353, p<0.05) and to increased OIL_TCD_ - PBMCe_TCD_ (0.58 p<0.05).Given that this has been identified as a regulator of antitumor immunity and a marker of lymphocyte exhaustion (*19*), these findings potentially suggest that the use of immune checkpoint inhibitors could further enhance OIL activity. We additionally note that all correlations were also maintained when considering the primary tumor types separately as appendiceal and mesothelioma primary tumors (Supplementary Figure S5).

### OILs Show Increased Polyfunctional Effector, Stimulatory, and Chemoattractive Activities

Because CD8^+^ effector cells are important for successful ACT therapy and because the abundance of these cells are seen to increase among OILs, we next sought to evaluate the polyfunctionality of the CD8^+^ fraction in OILs compared to that of PBMCes and TILs. The presence of polyfunctional effector immune cells (*i.e.,* cells producing multiple cytokines, chemokines, or cytotoxic granules simultaneously) has been shown to be critical to overall survival for many cancer patients (*20*),(*21*). Therefore, to determine if OIL CD8^+^ cells had increased polyfunctionality – and if so, in what functional cytokine groups they represent – cells derived from three patients with appendiceal cancer and one with mesothelioma were evaluated at the end of expansion phase (see Figure 1A). Measurements were conducted at this specific timepoint to determine their potential activity upon therapeutic administration. The CD8^+^ cells, which included cytotoxic T cells and some NK cells, were isolated independently from OILs, PBMCes, and TILs using magnetic activated cell sorting. Polyfunctionality was subsequently assessed using the Isoplexis IsoCode human single cell cytokine secretome analysis kit, which measures a preset panel of 32 multiplexed secreted cytokines specific to the human adaptive immune system pathway at a single-cell resolution for complete immune cell functional characterization (Figure 4A-B).

**Figure 4.**
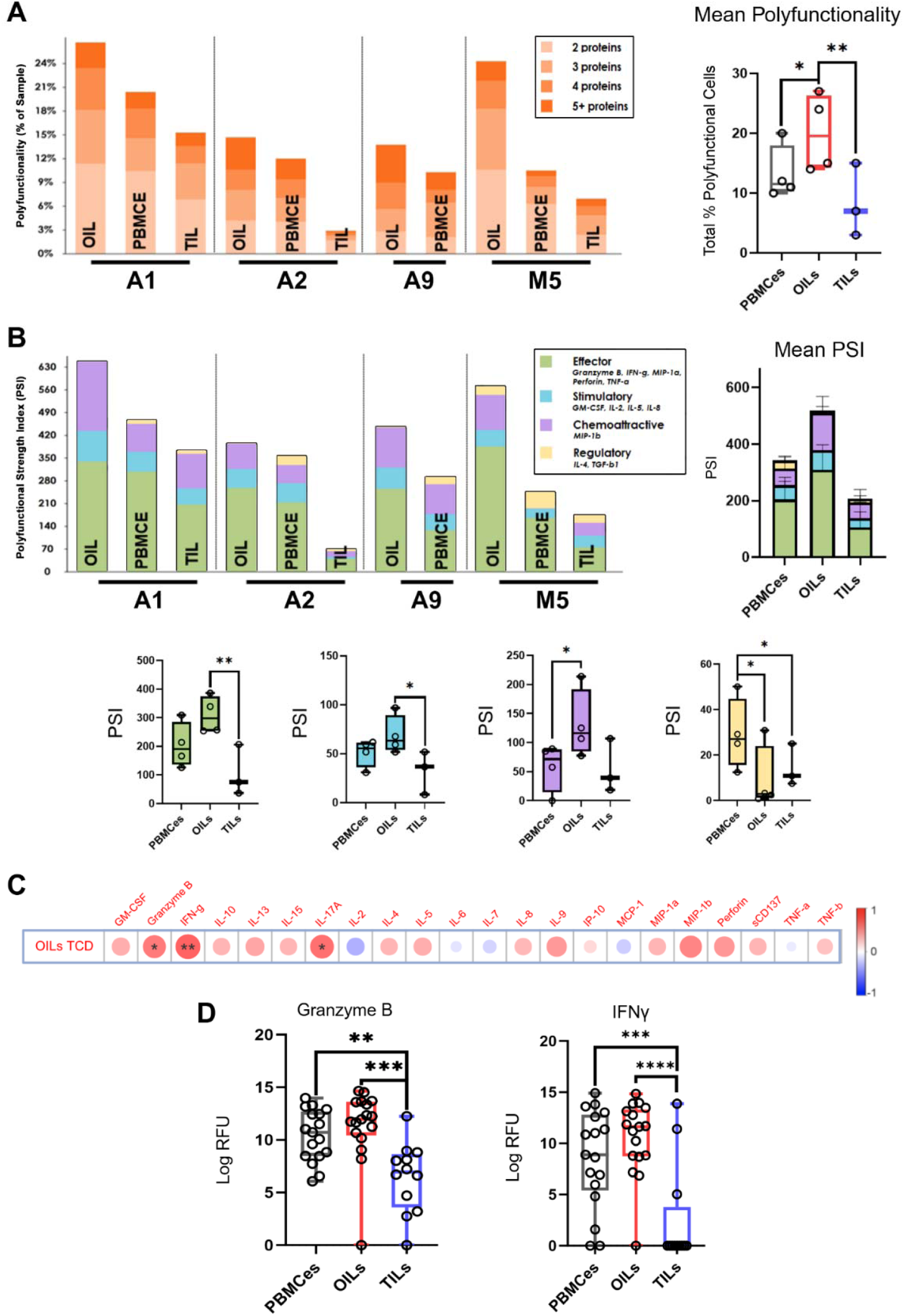
Proteomic analysis showing increased OIL polyfunctional cytotoxic phenotype. **(A)** Comparison of the percent of polyfunctional cells (multiple cytokine secretions per cell) in total CD8^+^ cell populations between immune cell groups (PBMCe), (OIL), and (TIL)] for 4 patients (n=3 appendiceal, n=1 mesothelioma) measured by Isoplexis Single Cell Secretome. The mean percent of total polyfunctional cells across all patients is also depicted. **(B)** Single cell polyfunctional strength index scores (PSI; number of single cells producing the cytokine multiplied by the cytokine signal strength) from the same patients in A to characterize the predominate cytokines produced. Cytokines are grouped into effector (green: Granzyme B, IFN-γ, MIP-1α, Perforin, TNF-α), stimulatory (blue: GM-CSF, IL-2, IL-5, IL-8), chemoattractive (purple: MIP-1β), and regulatory functions (yellow: IL-4, TGF-β1). Box plots of both total PSI scores and PSI scores for each cytokine group are also shown for the combined patient groups. **(C)** Correlation heatmap for cytokine expression in the cytotoxicity assay co-culture media [relative fluorescence units (RFU), measured by Isoplexis CodePlex analysis] in relation to OIL-induced tumor cell death (TCD) (n=17 patients; 13 appendiceal and 4 mesothelioma). **(D)** Individual expression of effector cytokines Granzyme B and INF-γ measured from the same co-culture media in (C). Data points in all box plots represent individual patients. Box plots present minimum, median, and maximum values. Statistical significance calculated for linear mixed models using post hoc Tukey method. Pearson correlation coefficients analyzed using T-test. *p<0.05 **p< 0.01 ***p< 0.001 ****p< 0.0001.

Across the 4 patients, OILs were found to have significantly more individual polyfunctional cells (*i.e.,* secreting 2 or more cytokines) than either TILs or PBMCe cells, indicating an overall increase in the presence of polyfunctional immune cells (OILs 20% vs PBMCe 13.3% p<0.05; and OILs 20% vs TILs 8.3% p 0.01; Figure 4A). When characterized by functional activity, OILs had a significantly increased polyfunctional strength index (PSI) of effector cytokine (Granzyme B, IFN-g, MIP-1a, Perforin, and TNF-a) when compared to TILs (OILs 309 vs TILs 106; p<0.01) (Figure 4B). In addition, OILs showed a significantly increased PSI for chemoattractive cytokines (MIP-1b) compared to PBMCe cells (OILs 131 vs PBMCe 58; p<0.05) as well as an increased PSI for stimulatory cytokines (GM-CSF, IL-2, IL-5, IL-8) compared to TILs (OILs 69 vs TILs 32; p<0.05). These findings highlighted the increased effector, stimulatory, and chemoattractive functions of OILs, as well as an overall enhancement in the number of OILs that could simultaneously perform these actions compared to PBMCe cells and TILs.

With the significant increase in OIL CD8^+^ cytokine signal secretion demonstrating the robust effector potential of these cells at the end of expansion phase, we next evaluated bulk cytokine production in co-culture media during the *in vitro* tumor cytotoxicity stage as a surrogate measurement of tumor-induced OIL activity post-treatment. To achieve this, bulk cytokine analyses were performed on co-culture media supernatant samples of 13 appendiceal patients and 4 mesothelioma patients using the Isoplexis Codeplex Adaptive Immune Panel, a predetermined multiplex assay of 22 target cytokines that play key roles in the adaptive immune system. For these patients, the relative fluorescent unit (RFU) of each cytokine measured in co-culture media was correlated with OIL_TCD_ (Figure 4C), similar to the data shown in Figure 3C. Increased OIL_TCD_ showed significant positive correlation with expression of several cytokines including the key stimulatory and effector molecules Granzyme B, IL-17a and IFNγ (Granzyme B 0.52, p<0.05; IFNγ 0.61, p<0.01; IL17 0.55, p<0.05). When comparing the RFUs of individual key effector cytokines expressed in OIL co-culture media to that of PBMCes and TILs (Figure 4D), OIL and PBMCe co-cultures expressed significantly increased cytotoxicity-associated cytokines over TILs when analyzing Granzyme B (OILs 11.3 vs TILs 6.55 p<0.01 and PBMCe 10.44 vs TILs p<0.01), and IFNγ (OILs 10.5 vs TILs 2.88 p<0.0001 and PBMCe 8.55 vs TILs p<0.001). All results were again preserved when analyzing the primary tumor types separately (Supplementary Fig. S6).

### Granzyme A Production is Consistently Upregulated during OIL-Induced Cytotoxicity

To further confirm the immune cell phenotypic profiles and activation status of PBMCes, OILs, and TILs, and to better explain the increased OIL_TCD_ through elucidation of potential apoptotic pathways within the tumor cells, co-cultures of tumor organoids with immune cells were analyzed by proteomic digital spatial profiling using nanoString GeoMX. Tumor organoids were cultured from the primary tumor cells of 5 mesothelioma patients, and matched PBMCes, OILs, or TILs were added to the organoid media to enable infiltration and interaction for 7 days. Through immunofluorescent imaging on day 7, we observed infiltration of CD45^+^ immune cells into the organoid as well as the presence of mesothelin^+^ tumor cells (Figure 5A). Regions of interest (ROI, white line) were selected for organoid co-cultures with each of the three immune cell groups. The ROIs were then segmented by cell type (CD45^+^ or mesothelin^+^) for regional analysis of protein expression using a specific immune-oncology panel. The mean z-scores of all mesothelin^+^ and CD45^+^ segments for each co-culture category were then compared using a heatmap (Figure 5B and C). The CD45+ segments in OIL co-cultures showed a relative increase in HLA-DR expression (OIL 1.04, PBMCe -0.08, TIL -0.96) that paralleled our prior flow cytometry analyses (see Supplementary Fig S4) as well as increased CD3 (OIL 0.71, PBMCe 0.44, TIL -1.14) and Granzyme A expression (OIL 1.12, PBMCe -0.32, TIL -0.79). The CD45^+^ segments from PBMCe co-cultures showed a relative increase in cleaved-caspase 9 expression, likely indicating some PBMCe death at the chosen experimental timepoint. Similar analysis of the mesothelin^+^ tumor segments from the OIL co-cultures showed a relative increase in antigen presentation (beta-2 microglobulin; B2M; OIL 0.99, PBMCe 0.02, TIL -1.01) and p53 expression (OIL 1.15, PBMCe -0.54, TIL -0.61).

**Figure 5.**
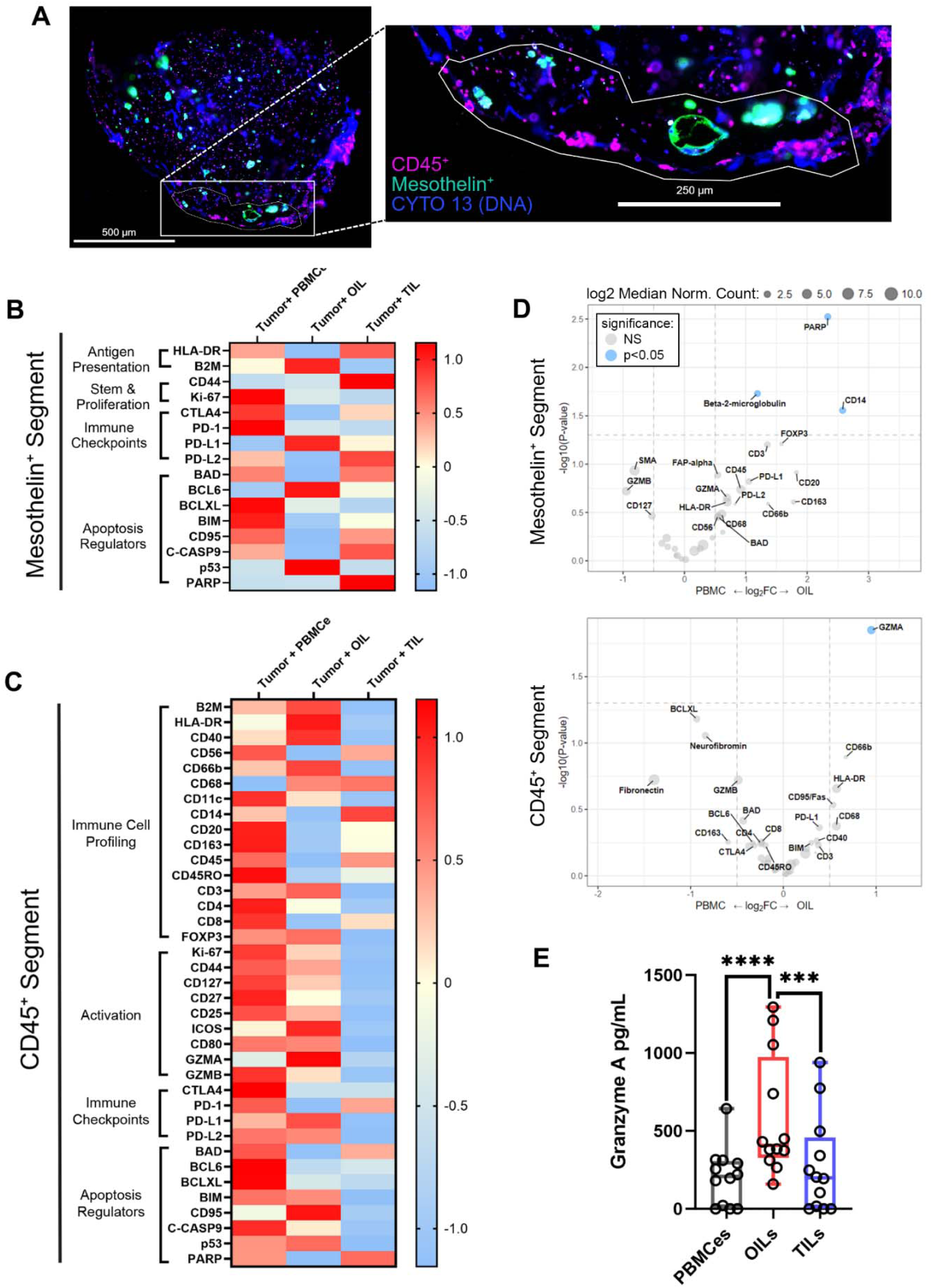
Spatial proteomic analysis of OILs-tumor interaction and OIL induced tumor cell death indicates increased Granzyme A activity in OILs. **(A)** Representative immunofluorescent image for GeoMX spatial proteomic analysis of mesothelioma tumor organoid (mesothelin^+^, cyan) co-cultured with OILs (CD45^+^, magenta), collected on day 7. Blue indicates nucleic acid (CYTO 13). Image on left is of the entire 5µl ECM dome, with the inset image on the right including a white line to indicate region of interest (ROI) selected for analysis. Within ROIs, CD45^+^ or mesothelin^+^ segmented areas of illumination were independently assessed for varying protein expression using the immuno-oncology panel from nanoString. For the experiment, 5 mesothelioma patients tumor organoids were cultured with PBMCes, OILs, or TILs. From each patient, 1-3 total ROIs were selected per immune-organoid co-culture condition, divided over 2-3 culture replicates. **(B)** Relative protein expression heatmap of combined mesothelin^+^ segments compared between co-culture groups. The color gradient represents expression levels as z-scores. **(C)** Relative protein expression heatmap of CD45^+^ tumor cell segments compared between co-culture groups. **(D)** Volcano plots comparing differential expressed proteins between OIL and PBMCe co-cultures for either CD45^+^ or mesothelin^+^ segments. Dashed lines indicate the significance threshold defined as log2 fold change >0.5 and blue indicates p<0.05. **(E)** Results of enzyme-linked immunosorbent assay to quantify Granzyme A production in PBMCe, OIL, and TIL co-culture media supernatant on culture day 7 (n=12 patients). Data points in box plots represent the means for individual patients, measured in triplicate technical assay replicates on a pooled culture media supernatant sample from at least 3 independent culture replicates. Box plots present minimum, median, and maximum values. Statistical significance calculated using linear mixed model to account for patient variability with post hoc Tukey analysis. ***p< 0.001 ****p< 0.0001.

Differential protein expression analysis specifically comparing mesothelin^+^ tumor segments between PBMCe and OIL co-cultures (Figure 5D, left) showed a significant increase in the fold change of B2M and poly(ADP-ribose)-polymerase (PARP-1) expression in mesothelin^+^ cells when tumor organoids were co-cultured with OILs (p<0.05). The increase in PARP-1 indicated an upregulation of DNA repair mechanisms but PARP1 hyperactivation has also been shown to results in non-caspase dependent cell death pathways(*22*). In the corresponding CD45^+^ cell segments for PBMCe and OIL co-cultures (Figure 5D, right), the fold change of differentially-expressed proteins highlighted a significant upregulation of Granzyme A expression in OIL co-cultures (p<0.05). Measurement of Granzyme A in the media from both appendiceal and mesothelioma organoid co-cultures confirmed these findings (mean OILs 587.7 pg/mL vs TILs 276.7 p<0.001; and OILs vs PBMCe 203.2 p<0.0001, Figure 5E), and Supplementary Figure S7. Overall, these results suggested that the significant upregulation of PARP-1 in the tumor cells co-cultured with OILs was likely related to the significant increase in Granzyme A activity and its capacity to initiate pathways of tumor cell death.

## Discussion

Herein, we have presented initial results from a novel method to generate cells potentially suitable for ACT therapy. Our system interfaces PBMCs – a readily available and renewable supply of autologous immune cells – with matched APCs and tumor cells in a microfluidic platform to create OILs: a population of lymphocytes with tumor-specific antigen priming. As a first demonstration of the approach, we employed specimens derived from patients with stage IV peritoneal cancers, including appendiceal and peritoneal mesothelioma malignancies. We showed that OILs significantly outperformed naturally-occurring TILs in induced primary tumor cell cytotoxicity *ex vivo* and determined that this behavior was also linked to increased polyfunctionality and secreted effector cytokines like Granzyme A. These metrics have been established previously as essential predictors of anti-tumor response and consequently of clinical efficacy. By testing OILS on patient-matched primary tumor tissue, we demonstrated that OILs exhibited strong effector function across a heterogenous population of samples, including activity that compared favorably to TILs in the subset of patients from which they could be retrieved.

Similar to the first FDA approved TIL therapy (Lifileucel), our approach leverages autologous lymphocytes and requires approximately 3 weeks to perform in its current form. Recent Lifileucel clinical trials showed an overall response rate of 36% across 66 patients and 31% across 153 patients with metastatic melanoma (*1, 23*). Failure of this therapy could have been due to either an inability to procure sufficient TILS from resected tissue or ineffective T cell priming *in situ (24)*. OILs address each of these important clinical challenges. While extant TIL treatment use irradiated, allogeneic peripheral blood cells to provide co-stimulation for lymphocyte proliferation, our methodology instead employed the tumor tissue itself, thereby exposing T cells to a diversity of tumor-specific antigens. Prior work (*25, 26*) has demonstrated the presence of TIL clonotypes in peripheral blood, suggesting that our use of PBMCs not only provided a renewable source of lymphocytes for priming but also a reservoir of naturally-activated cells. It is also important to mention that appendiceal cancers are generally considered non-immunogenic, limiting the applicability of TIL controls. Nonetheless, we have previously shown in a preclinical appendiceal cancer model that approximately 21% of appendiceal cancer organoids respond to Pembrolizumab with 11% responding to Nivolumab (*9*). While in the present study, OILs at the level of the entire cohort show a modest average increase over PBMCs (∼10% TCD), nearly half of patients (46%, 6/13) achieved an additional median cytotoxicity of 34.7% over PBMCe, with observed cytotoxicity ranging from 60.4% to 92.9%. This supports the potential of OILs for individualized therapy, identifying patients who could meaningfully benefit from this strategy.

To further explore the mechanisms of OILs efficacy, we also sought to perform phenotypic analyses across patients. While immune cell phenotype analysis alone could not fully explain the increased TCD of OILs, we found that the OILs showed conspicuous increase in CD8^+^ T cells. CD8^+^ T cells have previously been shown to play a prominent role in tumor cytotoxicity, with their abundances correlating with favorable clinical outcomes (*27, 28*). In correlation analysis of immune cell phenotype to OIL_TCD_, we also found a significant relationship between the presence of NK cells and increased OIL cytotoxicity. NK cell-mediated TCD, either direct or through the enhancement of T cell-mediated training and killing, is well documented (*29*),(*30*),(*31*),(*27*). While overall numbers of NK cells were low in the final OIL product, this cell population likely contributed to the success of the OIL therapeutic. The relative populations of immune cell phenotypes present among OILs also provided insight into why these cells displayed increased anti-tumor activity over TILs and their untrained PBMCe counterparts, and further in-depth functional analysis is warranted to better understand therapeutic response mediated by diverse interactions among lymphocyte populations.

Furthermore, we observed that for a small subset of patients (n=2) within the appendiceal cohort, *in vitro* static expansion of PBMCs resulted in TCDs that were comparable to or even greater than OILs, suggesting that further characterization of baseline PBMC populations was warranted. Phenotypic analyses of these samples indicated that high PBMCe activity correlated directly with the percentage of HLA-DR expression in the initial (PBMCb) T cells, particularly within the CD8^+^ T population where it is known to be expressed on late-stage activated cells (up to 5.8% of peripheral T cells (*32*)). HLA-DR on cytotoxic T cells has been strongly associated with positive response to neoadjuvant chemotherapy treatment in breast cancer patients (*14, 33*) and has been demonstrated as a highly-proliferative and anti-tumor cytotoxic effector phenotype in hepatocellular carcinoma (*32*). These properties suggest a pathway to pre-existing T cell activation *in vivo* and suggest that expansion of peripheral blood alone could yield a useable product for ACT, especially for PBMCs rich in HLA-DR. Further investigation may delineate a more precise immunophenotype linked to this response.

One important consideration in this study was patient-to-patient variability. For example, the diverse prior treatments experienced by the subjects (see Table 1) could have impacted circulating PBMCs prior to collection, including prior chemotherapy treatments. Given the heterogeneity of most solid tumors, it is also likely that lymphocyte populations were trained on different neoantigens from patient to patient. For this reason, clonality assessment like that used to characterize Lifileucel (*34*) was deemed unnecessary. In future research, a well-characterized tumor model could be used to assess the binding capacity of the tetramer to a known tumor antigen for validation of lymphocyte priming in our platform. Despite sample heterogeneity, the majority of patients across all peritoneal cancers studied here showed improved tumor cell death with OILs over TILs, thereby demonstrating the potential of our platform to generate tumor-specific immune cells and expand the pool of patients that are candidates for ACT therapy.

In conclusion, this study has demonstrated a training platform capable of using readily-available PBMCs to generate a robust, mixed-cell population of polyfunctional effector lymphocytes with tumor cytotoxic activity. Additional research will be required to fully understand the cellular mechanisms underlying the enhanced tumor-targeting capacity of OILs, including investigation into the CD8^+^ T cell and NK cell phenotypes and their roles in Granzyme A-induced TCD, potentially through investigation of mitochondria membrane potential disturbance and production of reactive oxygen species. Additionally, future extensions of the approach into animal models will enable the *in vivo* administration of OILs to be assessed as well, elucidating tumor-targeting capacity in the peritoneal cavity as a pre-clinical model for therapeutic use. Ultimately, OILs represent a new potential ACT modality using small tumor specimens collected during therapeutic cytoreductive surgery, or even through small core biopsies. With the potential to be a renewable therapeutic cell resource through repeated collection of patient PBMCs, we anticipate their utility in targeting a broad array of tumors that are currently untreatable with available therapies.

## Materials and Methods

### Tumor Tissue Collection and Processing

Between 2022 and 2024, abdominal tumors, peripheral blood, and normal lymph nodes or normal spleen were collected under IRB approved protocols (IRB00040474, IRB00043947), from 10 peritoneal mesothelioma and 21 appendiceal cancer with peritoneal metastasis patients undergoing cytoreductive surgeries. Table 1 describes the patient and tumor characteristics, as well as in which experiments each patient was included. For cases where spleen was obtained, these patients were already undergoing total splenectomy due to severity of tumor encasement around the splenic capsule. A small portion of unaffected spleen, as determined by pathologist, was sent for chip fabrication. Patients were included in all experiments possible when sufficient numbers of each cell population (tumor, PBMCb, PBMCe, OIL, and TILs) were available. Intraoperative specimen collection was performed by the operating surgeon from multiple spatially distinct sites of tumor involvement, in an attempt to increase representation of clonal neoantigens of the underlying malignancy within the harvested tumor sample. Specimens were maintained in RPMI-1640 media (Gibco) at 4 C and transferred within 2 hours by a dedicated procurement agent to the Wake Forest Organoid Research Center laboratory for tissue processing.

For tumor cell isolation, tumors samples, with weights ranging between 150 mg to 2 mg (data not shown), were rinsed twice in Dulbecco’s phosphate buffered saline (DPBS) with antibiotic cocktail (1x antibiotic-antimycotic, Gibco). Tumors and lymph/node spleen tissue were minced into ∼1 mm^3^ pieces and digested for 75 minutes with agitation at 37 °C in 3 mL/1 mg tissue of low glucose DMEM media containing 100,000 cytidine deaminase U/mL collagenase HA 200 (001-100, VitaCyte), 22,000 narcissus pseudonarcissus agglutinin U/mL BP Protease (003-1000, VitaCyte), and 200 U/ml DNase I (Sigma). Enzyme activity was quenched by adding an equal volume of cold RPMI media containing 10% fetal bovine serum (FBS). The dissociated cells were filtered through a sterile vacuum filtration kit with 60 µM pore size (Millipore) and pelleted by centrifugation. The cell pellet was incubated with 1x red blood cell lysis buffer (BD Biosciences) for 3 minutes. After quenching and washing with media, cells were resuspended in Organoid Media (advanced DMEM/F12 containing 5% FBS, 50 ng/mL epidermal growth factor (EGF), 1% Insulin-Transferrin-Selenium (ITS), 1% L-glutamine and 1x antibiotic-antimycotic). To isolate PBMCs, whole blood was processed using Ficoll-Paque PLUS and its protocol (GE Healthcare). All cells were counted using a NucleoCounter NC-200 (Chemometec). Tumor cells were either used directly in the microfluidic chip assembly, cryopreserved in Recovery Cell Culture Freezing Medium (12648010, Gibco) for downstream organoid fabrication for GeoMX analyses, or seeded onto Geltrex-coated tissue culture flasks for propagation for cytotoxicity analysis co-culture for flow cytometry. A portion of PBMCs (approximately 8-10 million cells) were cryopreserved in Recovery Cell Culture Freezing Medium to serve as the baseline PBMCs for flow cytometry analysis.

### Microfluidic Chip Assembly

Microfluidic chips were assembled from the following layers: 1) a plain glass slide (12-550-A3; Fisherbrand), 2) 0.074 mm thick double-sided adherent membrane (DST, 3M162880-ND; Digikey), 3) 1.5 mm thick poly-methyl methacrylate (PMMA; 8560K239, McMaster-Carr) 4) another DST layer, and 5) a final 3 mm thick PMMA layer. The first DST layer was cut with entry and exit holes on either side of an open chamber space using a laser cutter (SKU MUSE3D-45W, Full Spectrum Laser). The two PMMA layers and an additional DST layer were cut with matching entry and exit holes to the chamber tape layer. The chip was assembled by aligning and stacking glass side, tape layer with open chamber space, PMMA layer, tape layer, and final PMMA layer (Figure 1). The immune-enhanced patient-derived tumor organoid (iPTO) chip was prepared by mixing 1.5x10^6 lymph node or spleen cells with 5x10^5 tumor cells and then seeding in a basement membrane hydrogel. In the initial generation of chips (patients A1-A11, A21, and M1-M7), this tumor-secondary lymphoid tissue cell combination was seeded within a HyStem-HP hydrogel matrix (heprasil:gelin:extralink in 2:2:1 ratio; GS1006F, Advanced Biomatrix) that was crosslinked by exposure to ultraviolet (UV) light (365 nm, 18W/cm2) at a distance of 4 inches for 7 seconds with the light probe (BlueWave 75 V.2 UV spot lamp; Dymax Corp., Torrington, CT), which created individual organoid columns within the chip. Chips were flushed with Dulbecco’s Phosphate-Buffered Saline (DPBS; Gibco) to remove excess hydrogel between the crosslinked organoid columns. To increase exposure of immune to tumor cells, a second generation of chips (patients A12-A20 and M8-M10) were assembled using similar layers as above, but substituting a cell culture treated glass slide (1 well chamber slide, 177372PK; Lab-Tek) for the bottom layer that was pre-coated with Geltrex basement membrane material (A1569601; ThermoFisher Scientific) prior to cell seeding. Tumor and secondary lymphoid tissue cells were allowed to adhere to the Geltrex surface in the incubator overnight, prior to remainder of chip assembly, as described above. Polytetrafluorethylene (PTFE) transfer tubing (50-634-313, Masterflex) connected to a 15mL conical and peristaltic pump tubing (PT-2100PS-F, MPP) was inserted into the entry and exit holes of the chip, and the interface between tubing and holes was sealed by Epoxy (14270; Devcon).

### OIL Generation

To generate Organoid Interacting Lymphocytes (OILs), 2.0-4.0x10^6 PBMCs were added to the reservoirs in a 3mL mixture of 1:2 Organoid Media to T cell expansion media (10981, Stem Cell Technologies) supplemented with 5 ng/mL human IL-7 (200-07, Peprotech), 40 ng/mL human IL-2 (200-02, Peprotech), and 25 uL/mL anti-CD3/anti-CD28 antibodies mixture (10991, Stem Cell Technologies). Chips were then placed in a cell culture incubator with perfusion pump tubing attached to MP2 micro peristaltic pump (MP2-6-PC; Meinhard). Pump flow rate was set to 0.1 revolution/min (3 ul/min). After 4 days, an additional 3 mLs of media was added to the chip. Circulated PBMCs, now referred to as Organoid Interacting Lymphocytes (OILs) were collected after 7 days of continuous perfusion. Statistical analysis with linear mixed model comparing between OILs derived from the two different chip basement membrane materials indicated no differences between Geltrex or HyStem-HP hydrogel matrix derived OILs to induce tumor cell death in the final cytotoxicity assay (data not shown).

### PBMCe Generation

For uncirculated, static expanded PBMC controls (PBMCes), 1.0x10^6 cells were added to a culture flask in the same media as OILs for expansion over the same timeline. Following 7 days of circulation or flask expansion, both OILs and PBMCes were harvested and expanded for an additional 7 days in T cell expansion media (10981, Stem Cell Technologies) supplemented with 40 ng/mL human IL-2 (200-02, Peprotech), and 25 uL/mL anti-CD3/anti-CD28 antibodies mixture (10991, Stem Cell Technologies). At the end of cell expansion, individual patient live cell counts of OILs ranged from 5 million to 306 million (data not shown). A portion of OILs were also analyzed by flow cytometry at the end of expansion to quantify expression of tumor cell specific markers in the live, single cell population (appendiceal: CK20 Novus NP242616A94 and EpCAM eBioscience 11-5791-80; mesothelioma: mesothelin Abcam ab196235; Figure S8). Analysis revealed less than 1% expression of tumor cell markers in the total live OIL cells (appendiceal mean 0.13% and mesothelioma mean 0.92%).

### TIL Isolation

Isolation of tumor infiltrating lymphocytes (TILs) was performed following published methods on TIL development for clinical use. A portion of tumor tissue from the same surgical biopsies above (approximately 300 mg) was chopped into 2 mm fragments and cultured for 7 days in 24-well tissue culture plates in RPMI medium supplemented with 10% FBS and 6000 U/ml IL-2, as previously described(*35*). TILs were then collected and expanded in flask for another 7 days in complete RPMI medium with 6000 U/ml IL-2. Following the expansion phase, OILs, PBMCEs, and TILs were either used immediately for downstream analysis (cytotoxicity assay, flow cytometry analysis, and organoid co-cultures), or cryopreserved in Recovery Cell Culture Freezing Medium (12648010, Gibco) for Single Cell Secretome Analysis.

### Cytotoxicity/Tumor Cell Death Assay

Fresh PBMCes, OILs, and TILs were co-cultured with their patient matched tumor cells (passage 3) to determine induced tumor cell cytotoxicity. Herein, 5.0x10^4 tumor cells were seeded on tissue culture treated and Geltrex pre-coated 24 well plates for a 24-hours in Organoid Media. Media (containing any dead or non-adherent cells) was then removed after 24 hours and 2.5x10^5 PBMCes, OILs, or TILs were added in 1:1 Organoid Media to T cell Expansion Media supplemented with 40 nng/mL IL-2 for 24 hours, in triplicate. Tumor only control cultures were also maintained to measure spontaneous tumor cell death. Media supernatant was then collected and frozen at -80 C for Isoplexis CodePlex analysis. Total cells (tumor and immune) were trypsinized and collected for flow cytometry analysis.

### Flow Cytometry Analysis

For detection of PBMCe, OIL, and TIL cytotoxicity, trypsinized cells (both tumor and immune) were collected, washed, and then stained with 1:1000 Fixable Viability Stain (562247, BD Biosciences) in DPBS for 15 minutes at room temperature. Cells were then washed and incubated in 2% fetal bovine serum in DPBS containing antibody cocktail for cell-surface markers for 30 minutes on ice (Table S1), then washed again. Cells were then fixed in a 1% paraformaldehyde (PFA) solution and stored at 4 degrees until analysis on the flow cytometer (24-48 hours post-staining).

For the immune cell phenotype analysis of PBMCbs, PBMCes, OILs, and TILs, the PBMCbs were thawed and allowed to rest overnight in RPMI-1640 with 10% FBS, prior to analysis. The next day, all cells were stained in a similar manner to above with both fixable viability stain followed by an antibody cocktail for cell-surface markers (Table S2). Cells were then treated with Intracellular Fixation & Permeabilization Buffer kit (88-8824-00, eBioscience) according to manufacturer protocol. After a subsequent washing period, intracellular antibody cocktail (Table 4; FOXP3 only) was then added to each well at room temperature for 30 minutes. These cells were then washed and suspended in a 1% PFA solution before being refrigerated and analyzed by flow cytometry. Samples were collected using BDX20 Fortessa Flow Cytometer and analyzed using FACSexpress software.

### Single Cell Cytokine Secretome Analysis

Cryopreserved PBMCes, TILs, and OILs were prepared for the Isoplexis IsoCode human adaptive immune panel per manufacturer protocol (PRO-17 REV 1.0, Bruker Corporation). Briefly, cells were recovered overnight at 37 C in 5% CO_2_ incubator in RPMI medium supplemented with 10% FBS and 100U/ml IL-2. CD8^+^ T cells were isolated using human CD8 microbeads kit (Miltenyi Biotec, USA) according to manufacturer’s protocol. CD8^+^ T cells were then stimulated for 1.5 hours by brefeldin A-free PMA/ionomycin cell activation cocktail (BioLegend, USA) at 1.0x10^6^/ml cells in 48-well plate. Stimulated cells were stained with cell stain 405 (Bruker Corporation). Cells were washed and resuspended in complete RPMI at 1.0x10^6/ml cells, and then 30 uL of cells (3.0x10^4cells) were loaded into the IsoCode chips. Functional profiling (cytokine secretion) by single CD8+ cells was analyzed by the IsoPlexis Isolight instrument and IsoSpeak software (Bruker Corporation).

### Digital Spatial Proteomic Profiling of Organoid Co-Cultures

Patient-derived tumor organoids (PTOs) from the same tumor biopsies were generated by suspending tumor cells at 10 × 10^6^ cells/mL in a thiol-modified hyaluronan/heparin (Heprasil; Advanced Biomatrix) and methacrylated collagen (PhotoCol, Advanced BioMatrix, Carlsbad, CA) hydrogel in a 1:3 ratio. The cell/hydrogel mixture were placed into the wells of a 48-well non-tissue culture treated plate at 5 µL/well and followed by exposure for one second per organoid to UV light from a BlueWave 75 V.2 UV spot lamp (Dymax Corp., Torrington, CT). The organoids were then cultured for 7 days with Organoid Media. Co-culture was initiated by adding OILs, PBMCEs, and TILs at 5:1 of immune to tumor cell ratio and maintained in a media ratio of 1:2 Organoid Media to T cell Expansion Media supplemented with 0.1 ng/mL human IL-2 for 7 days. After, co-cultured organoids were collected for Nanostring GeoMX spatial analysis of protein expression, performed by the Wake Forest University School of Medicine Comparative Pathology Laboratory Core Facility. Briefly, organoids were fixed in 4% PFA, embedded in paraffin, sectioned, and pre-stained with anti-CD45 antibody (NanoString Tumor Morphology Kit) for hematopoietic immune cells (OILs, TILs, or PBMCs) and anti-mesothelin antibody for mesothelioma tumor cells (c). Using the nanoString GeoMX Immuno-Oncology Protein Panel kits, probes were hybridized to the slides according to manufacturer protocols. The geometric regions of interest (ROIs) were created around portions of the organoids enriched in CD45^+^ and mesothelin^+^ cells. These ROIs were then further segmented to independently analyze CD45^+^ and mesothelin^+^ cells for spatial protein expression in the different organoid co-culture conditions. Collected oligonucleotide barcodes were analyzed on the nanoString nCounter system. Quality control, normalization of data, and differential expression/statistical analysis were performed by data specialists at nanoString.

### Cytokine Profile Detection

Cytokine profiles of the tumor and immune cell co-culture media were determined using a CodePlex Secretome Human Adaptive Immune kit according to manufacturer’s protocol (PRO-11 REV 1.0, Bruker Corporation). Briefly, frozen media supernatant from the 24-hour cytotoxicity assay, pooled from 3 culture replicates, and CodePlex chips were completely thawed at room temperature. Pooled media samples and background controls were loaded in duplicate into the CodePlex chip at 5.5 μL/well. Chips were run under the CodePlex protocol on the IsoLight machine. Protein concentrations were analyzed by IsoSpeak software and data was presented as relative fluorescence unit (RFU). Additionally, granzyme A secretion in day 7 co-culture media from the organoids used for nanoString GeoMX analysis was quantified by sandwich enzyme-linked immunosorbent assay (ELISA) analysis using the Granzyme A ELISA kit (NB120238, Novus Biologicals) according to manufacturer’s protocol. Culture media supernatant samples were pooled from 3 culture replicates, and the pooled media was run in triplicate in the ELISA assay.

### Statistical Analysis

Data were analyzed using GraphPad Prism 10.1.2 software and R package. The statistical differences between treatment groups were determined by two-tailed, unpaired student’s *t*-test with unequal variance, linear mixed model analysis, or Pearson correlation coefficient, as indicated, with p<0.05 was considered significant. Normality was assessed using D’Agostino and Pearson test, and data were presented as means ± standard deviation or box plots with median, minimum, maximum, and lower/upper quartiles.

#### nanoString GeoMx Analysis

Data quality control and normalization were performed according to manufacturer guidelines. Linear mixed-effect models to account for sampling of multiple areas of illumination per patient per organoid co-culture were used to calculate differential protein expression on a per-protein basis from their normalized expression.

#### Normalization

In the case of cytotoxicity analysis, percent of dead cancer cells was normalized to spontaneous cell death using the linear transformation equation (*36*):

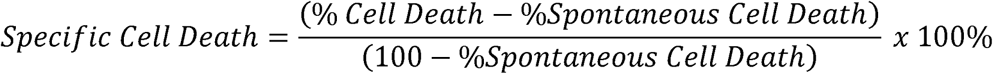

Spontaneous cell death refers to the mean dead cancer cell percent of 3 replicate tumor samples receiving no treatment.

#### Mixed Model Analysis

A linear mixed-effects model was applied to analyze the response variable, accounting for both fixed and random effects. The model was constructed using the lmer() function from the lme4 package with the following formula:

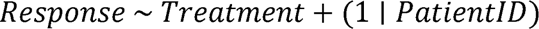

Here, treatment (i.e. PBMCb, PBMCe, OILs, or TILs) was considered a fixed effect, while Patient ID (e.g. A18) was treated as a random effect to account for inter-subject variability. The model’s summary, including estimates, standard errors, and p-values, was generated using the summary() function. Cytotoxicity analysis of mean appendiceal and mean mesothelioma treatment response was calculated by repeating the patient ID on the leftmost side for each of the 3 biological co-culture replicates examined. Other analyses using mixed effect model generation such as Grazyme A ELISA comparison used 1 Patient ID per treatment since culture replicate samples were pooled in those analyses.

#### Post Hoc Analysis

post hoc analysis was conducted using the emmeans() function from the emmeans package to identify significant difference among treatments after linear mixed effect model generation. This function computed estimated marginal means (EMMs) for the treatments, controlling for multiple comparisons using the Tukey method. Pairwise comparisons between treatment groups were then performed using the pairs() function, and their summary was obtained via the summary() function.

#### Correlation Analysis

Pearson correlation coefficients were calculated for the variables in the data frame. The correlation matrix and the associated p-values were computed using the cor.mtest function from the corrplot package, which performs a student t-test for each Pearson correlation coefficient. This function calculates both the correlation coefficients and the significance levels of the correlations. The corrplot() function (also from the corrplot package) was used to visualize the correlation matrix. The plot included: Correlation coefficients represented by colored circles.

#### Receiver Operating Characteristic Analysis

Receiver Operating Characteristic (ROC) analysis was used to identify the optimal cutoff point to determine which threshold would be consistent to predict increased TCD activity of PBMCes over OILs. Predictor variable comprised of baseline PBMC HLA-DR^+^ percentage in CD45^+^ cells. Response Variable was simplified to binary options whereby if PBMCes had higher induced cytotoxicity than OILs they were labeled 0 and if the reverse occurred outcomes were measured as 1. The ROC curve was generated using the ROC function from the pROC package in R. The ROC curve was used to calculate Youden’s Index, which is defined as sensitivity + specificity - 1. The optimal cutoff value was determined by identifying the threshold that maximized Youden’s Index.

## Acknowledgments

The authors wish to acknowledge the support of the Atrium Health Wake Forest Baptist Comprehensive Cancer Center Tumor Tissue and Pathology Shared Resource and Comparative Pathology Laboratory, including chief procurement agent Libby McWilliams, as well as Dr. David Caudell and Lisa O’Donnell for their support in the digital spatial profiling. This resource is supported by the National Cancer Institute’s Cancer Center Support Grant award number P30CA012197.

## Author contributions

CRS and DCH equally contributed to experimental design, data acquisition, data analysis, data visualization, original manuscript drafting, and manuscript review/editing. TL contributed to data acquisition, data analysis, and original manuscript drafting. MK contributed to experimental design and data acquisition. CJW and NE contributed to data analysis and data visualization. NW contributed to experimental design and data acquisition. SDF and RG contributed to experimental design and data acquisition. EAL and PS contributed to experimental design and sample collection. PT and LDM contributed to conceptualization, supervision, and manuscript review/editing. ARH, SS, and KIV equally contributed to conceptualization, supervision, data analysis and visualization, funding acquisition, and manuscript review/editing. All authors reviewed and approved the final manuscript.

## Data and materials availability

The data can be provided by Wake Forest University Health Sciences pending scientific review and a completed material transfer agreement. Requests for the data should be submitted to: Konstantinos Votanopoulos.

