## Supplemental Figures and Tables for "Enhancing Patient Lymphocyte Response to Peritoneal Malignancies Using a Personalized Immunocompetent Microfluidic Co-Culture Platform"

Cecilia R. Schaaf *et al.*

* *Adam R. Hall, Shay Soker, Konstantinos I. Votanopoulos*

**This file includes:**

Figs. S1 to S8

Tables S1 to S2

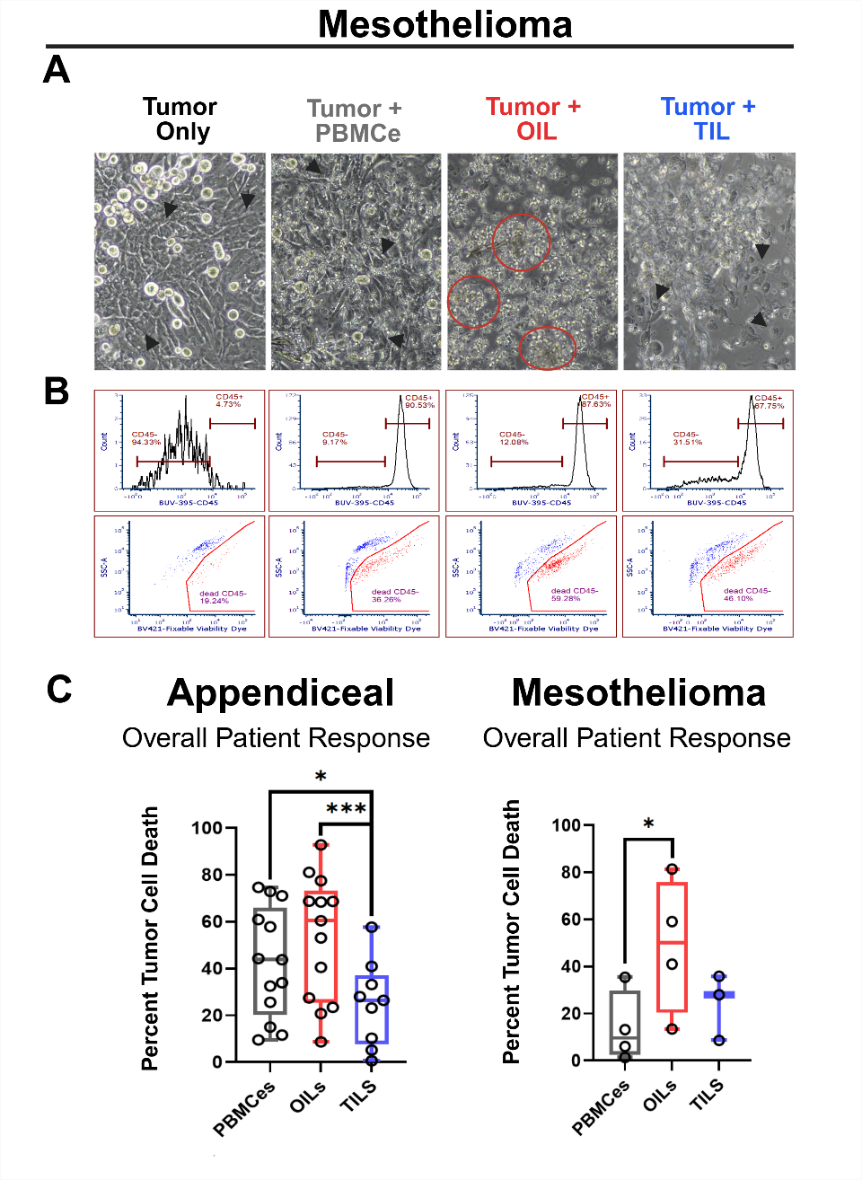

Fig. S1. Independent analysis of appendiceal and mesothelioma tumor types confirm treatment of peritoneal tumors with OILs resulted in increased tumor cell death. (A) Representative images from a typical patient (mesothelioma patient M8) showing tumor cells alone or co-cultured with immune cells PBMCe, OIL, and TIL following a 24-hour incubation to assess tumor cell death (TCD). Black arrows indicate tumor cells, red circles indicate immune cell clusters. (B) Representative flow cytometry gating strategy from the same patient in (A) to quantify death of tumor (CD45^-^ in histogram) cells alone or co-cultured with indicated immune cells via viability dye (scatterplot). (C) Assessment of overall patient response divided into appendiceal (left) and mesothelioma (right) cases (same data from Figure 2 in the main text) shown as the percent of TCD in co-cultures of tumor cells with PBMCes (gray), OILs (red), or TILs (blue). The percent of treatment-induced TCD was normalized to spontaneous cell death measured in a tumor-only control. Boxplots show the overall patient response (n=13 appendiceal, 4 mesothelioma, data points represent the mean of triplicate cultures measured for each patient). Data were compared using linear mixed models and post hoc Tukey method. *p< 0.05 **p< 0.01 ***p< 0.001.

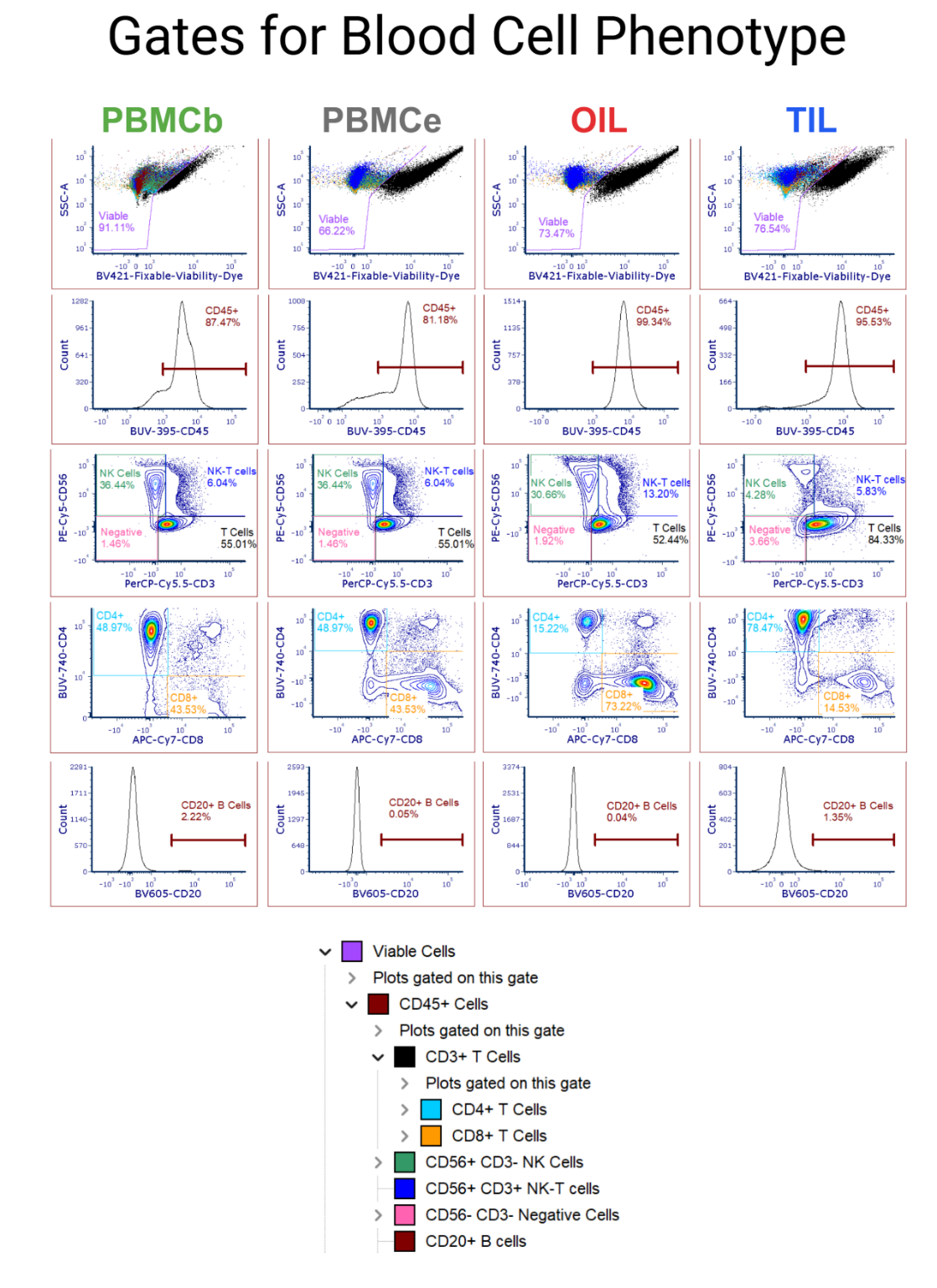

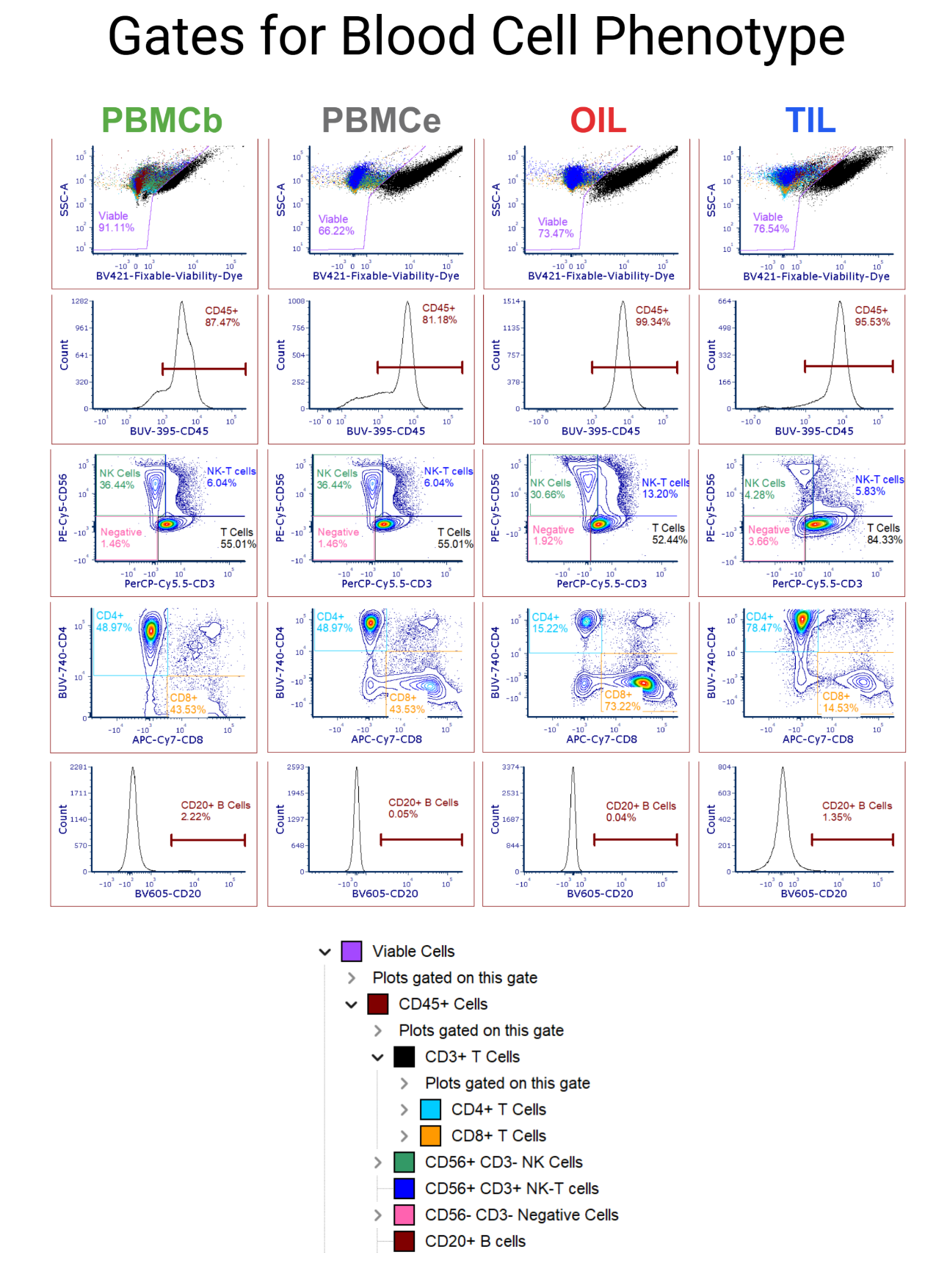

Fig. S2. Flow cytometry gating strategy to identify key immune cell phenotypes. Flow cytometry data for a representative patient, with each immune cell category (PBMCb, PBMCe, OIL, and TIL), gated on live, single cells. These were subsequently gated for CD45^+^ cells, which were further divided into CD3^+^, CD56^+^, and CD20^+^ populations. CD3^+^ cells were further identified into CD4^+^ and CD8^+^ populations. CD20^+^ cells were gated from the CD3^-^CD56^-^ population.

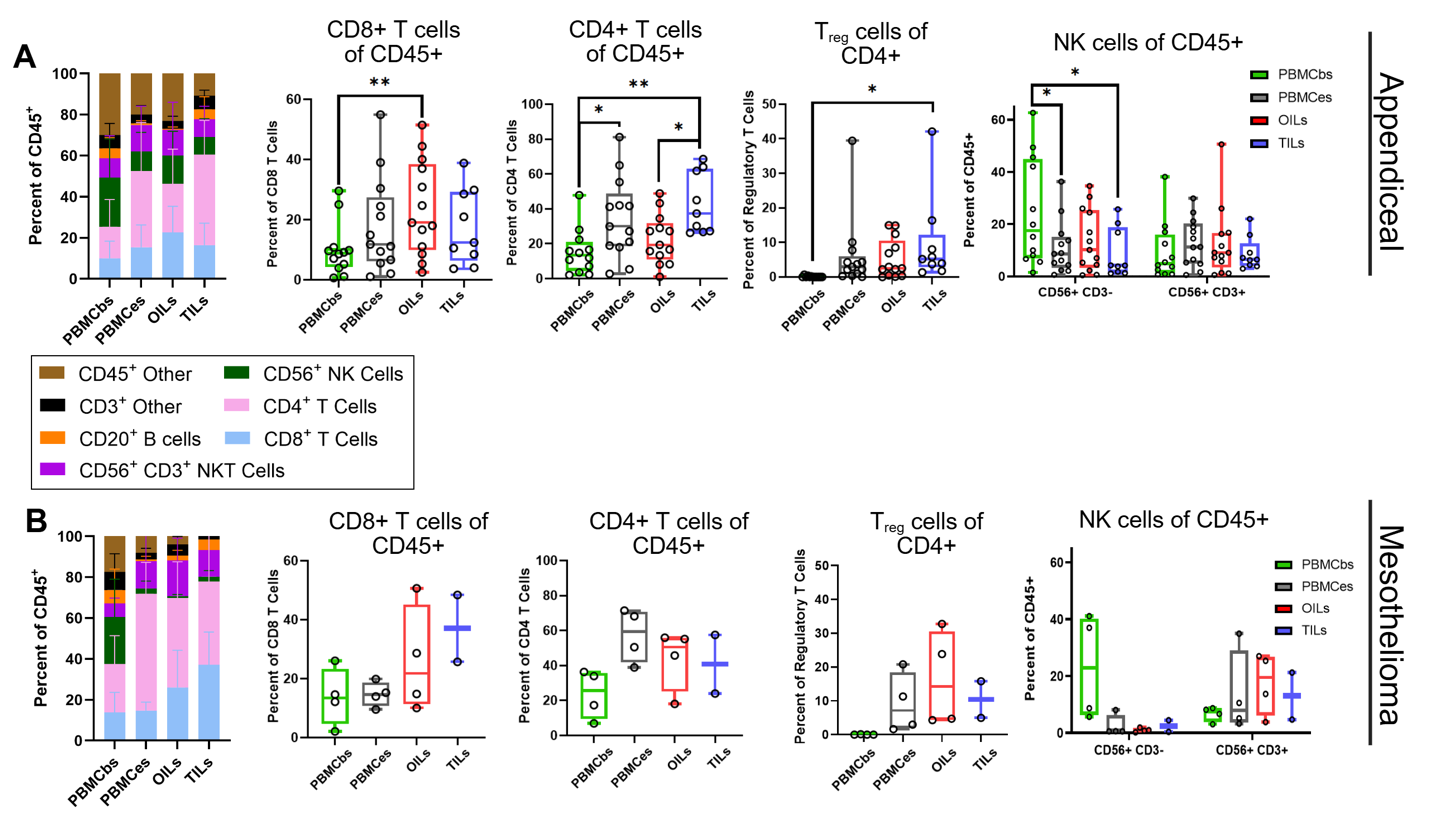

Fig. S3. Immune cell population phenotypes present in OILs are similar between appendiceal and mesothelioma primary tumor cases. Bar graphs (left) and box plots (right) showing the distribution of key immune cell phenotypes present among immune cell groups (PBMCb, PBMCe, OIL, and TIL) as determined by flow cytometry. The same peritoneal malignancy cases shown in Fig. 3a in the main text are divided into appendiceal (A) and mesothelioma (B) primary tumors. Data points in box plots represent individual patients (n=12 appendiceal in A and 4 mesothelioma in B). Each box plot shows minimum, median, and maximum values. Statistical significance calculated using linear mixed models and post hoc Tukey method. *p<0.05 **p<0.01.

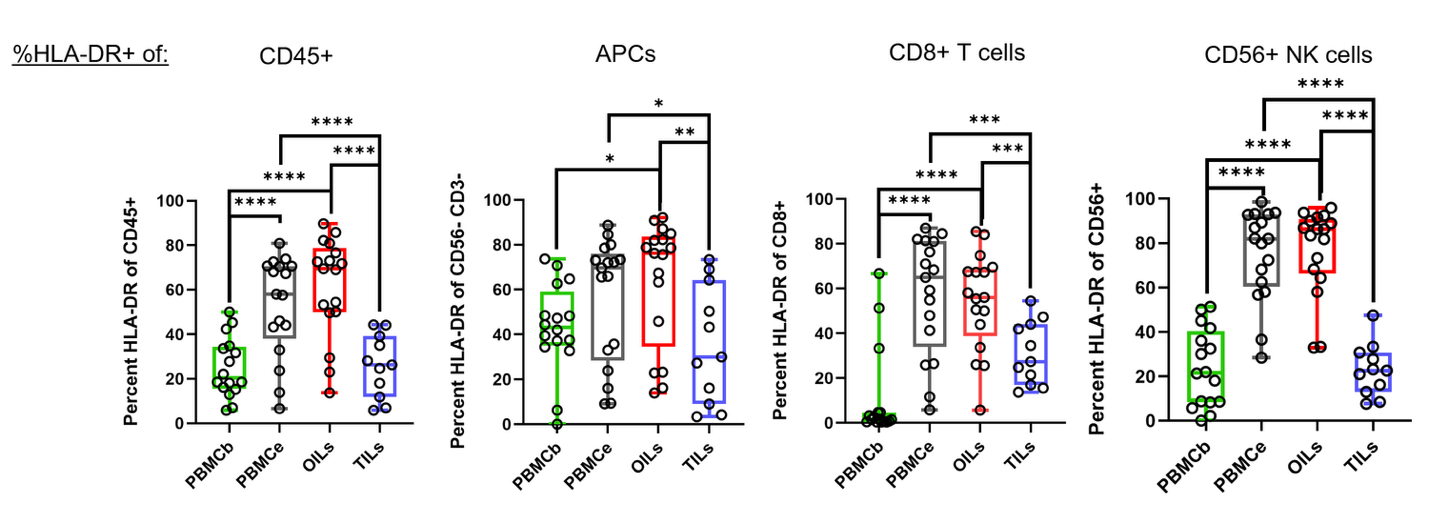

**Fig. S4. HLA-DR expression is increased in both PBMCe and OIL populations, indicating increased activation.** HLA-DR expression in patient-derived specimens determined via flow cytometry and displayed as a percentage of (*l-r*) total CD45^+^ cells, CD3^-^CD56^-^CD45^+^ Antigen Presenting Cells (APCs), CD8^+^ T cells, and CD56^+^ NK cells. Data points show values for all (n=16) peritoneal malignancy patient specimens. Each box plot shows minimum, median, and maximum values. Statistical significance calculated using linear mixed models and post hoc Tukey method. *p<0.05 **p<0.01 ***p<0.001 ****p<0 .0001.

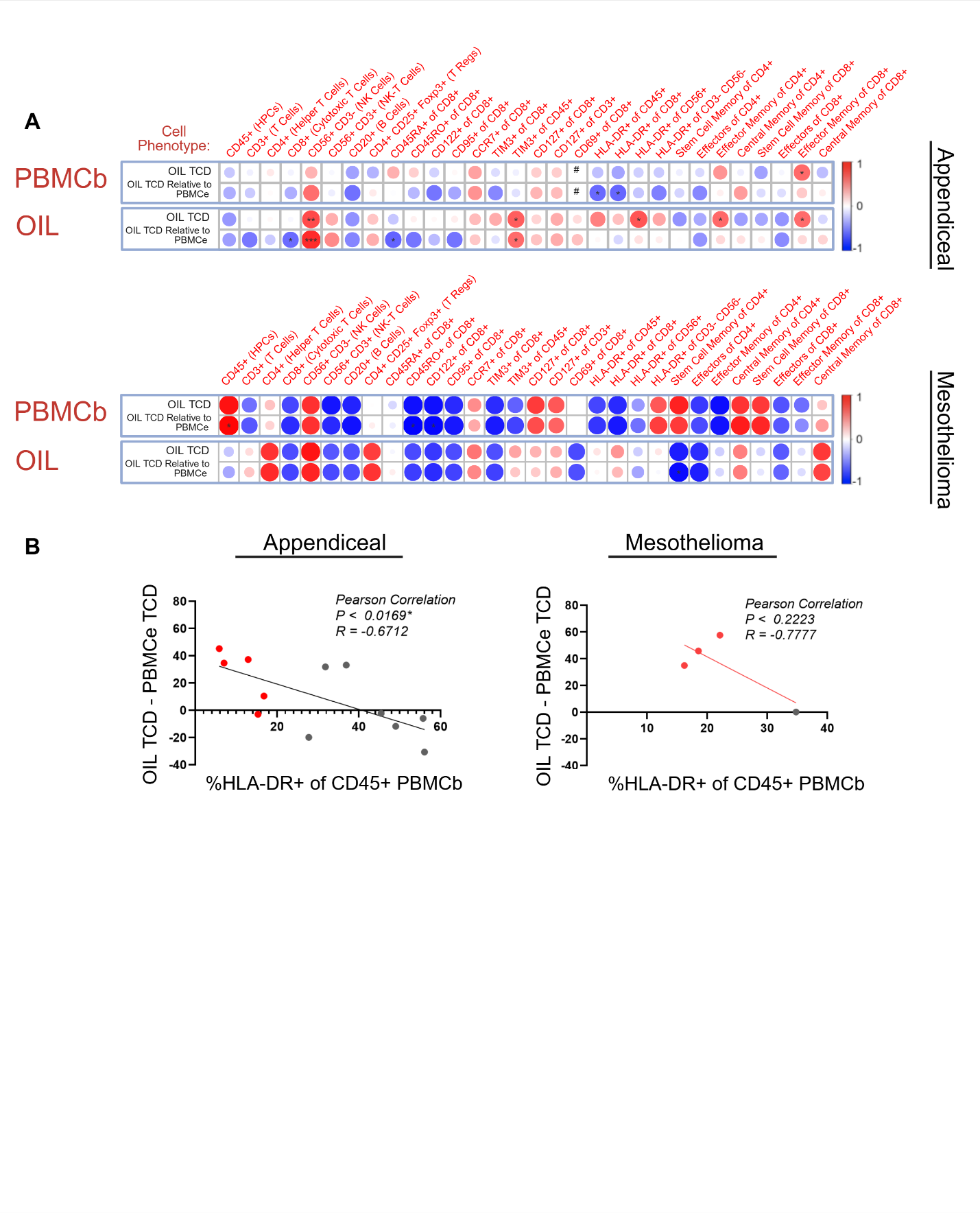

**Fig. S5. Immune cell population phenotypes analyzed independently by primary tumor type show negative correlation between OIL activity and baseline HLA-DR expression. (A)** Correlation heatmaps between TCD and PBMCb (upper data, red row title) or OIL (lower data, red row title) phenotypes, divided into appendiceal (top) and mesothelioma (bottom) primary tumors. Each data set depicts the normalized percent TCD for OILs (above) as well as the difference between OIL TCD and PBMCe TCD (below) as derived from Figure 2 in the main text. Expression of immune cell phenotype markers (red column labels) measured by flow cytometry are compared by Pearson correlation coefficient. The size and color of the heatmap indicate the Pearson correlation coefficient for each relationship, with a large, dark red circle indicating a perfect positive correlation equal to 1 and a large, dark blue circle indicating a perfect negative correlation equal to -1. * indicates a significance (*p<0.05; **p<0.01; ***p<0.001; **** p<0 .0001) determined by T-test while # indicates value not measured. **(B)** Pearson correlation plot comparing the percent of HLA-DR^+^CD45^+^ cells in the PBMCb population to the difference between OIL and PBMCe TCD, separated by low (red) versus high (gray) percent HLA-DR^+^CD45^+^ PBMCbs in appendiceal (left) and mesothelioma (right) primary tumors.

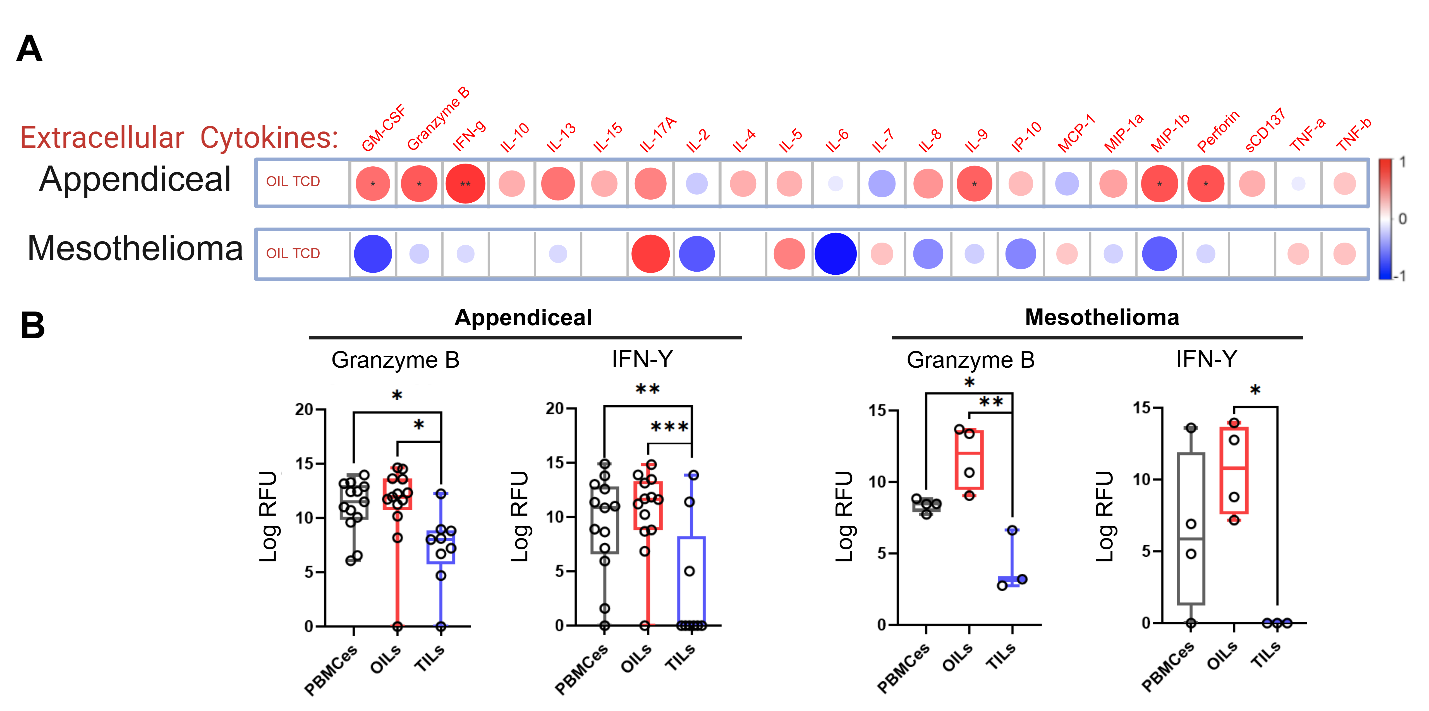

**Fig. S6. Independent proteomic analysis of appendiceal and mesothelioma primary tumors also demonstrate increased correlation in cytotoxicity associated cytokine expression in OIL effector function. (A)** Heatmap correlating expression of cytokines (red column labels) measured in co-culture media of the cytotoxicity assay [relative fluorescence units (RFU), measured by Isoplexis CodePlex analysis] to OIL-induced TCD (n=13 appendiceal and 4 mesothelioma). **(B)** Expression of effector cytokines Granzyme B and INF-γ measured from co-culture media with PBMCe, OIL, and TIL, divided into appendiceal (left) and mesothelioma (right) primary tumors. Data points represent individual patients. Box plots show minimum, median, and maximum values with error bars representing standard deviation. Statistical significance calculated using linear mixed models and post hoc Tukey method. Pearson correlation coefficients analyzed using T-test. *p<0.05 **p< 0.01 ***p< 0.001 ****p< 0.0001.

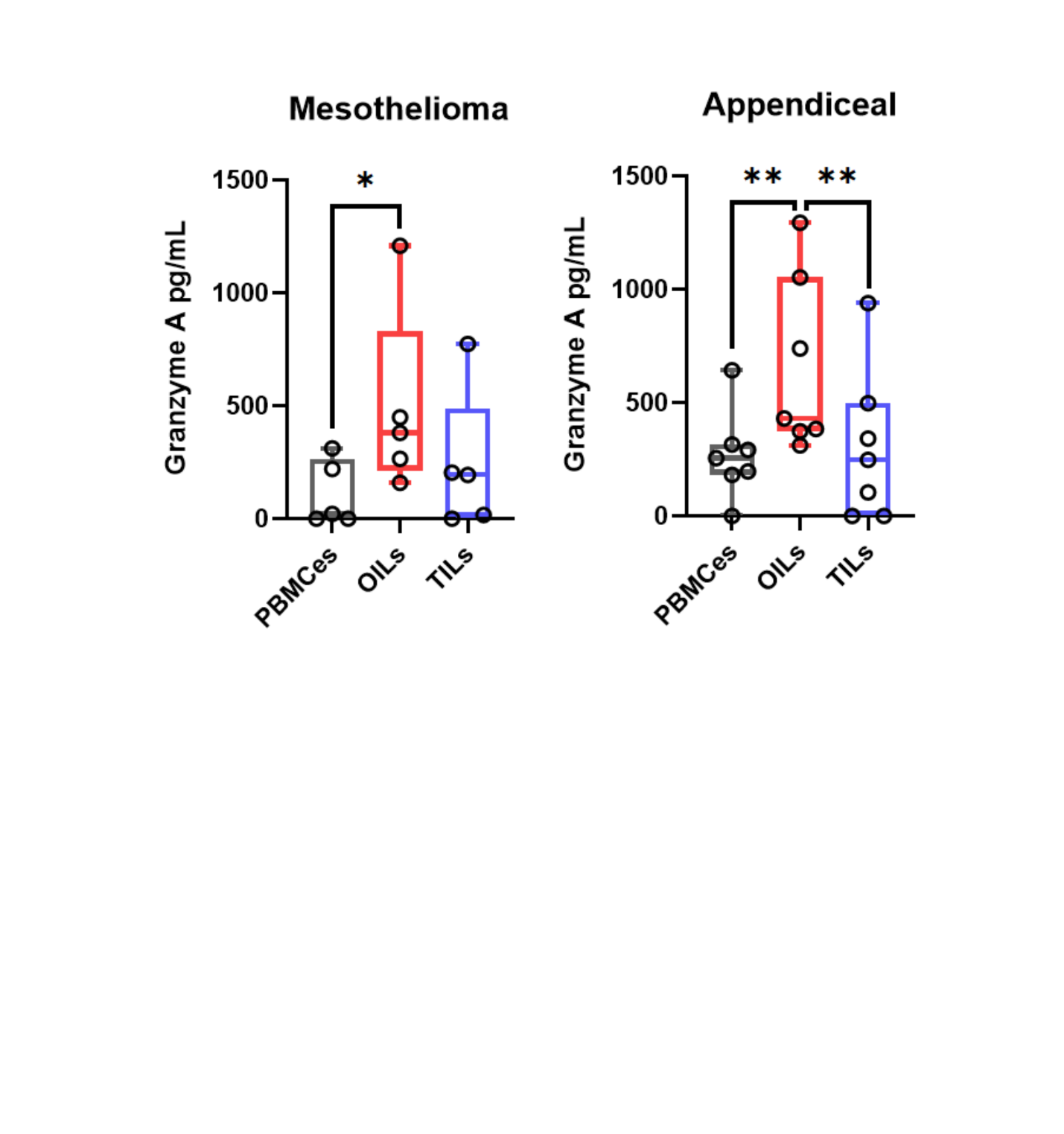

**Fig. S7. Independent analysis of mesothelioma and appendiceal tumor co-culture samples confirm increased Granzyme A activity in OILs.** Results of enzyme-linked immunosorbent assay to quantify Granzyme A production in PBMCe, OIL, and TIL co-culture media supernatant on culture day 7, divided into mesothelioma (n=5) and appendiceal (n=7) primary tumors. Data points represent individual patient specimens, measured in triplicate. Box plots show minimum, median, and maximum values with error bars representing standard deviation. Statistical significance calculated using linear mixed models and post hoc Tukey method. *p< 0.05 **p< 0.01.

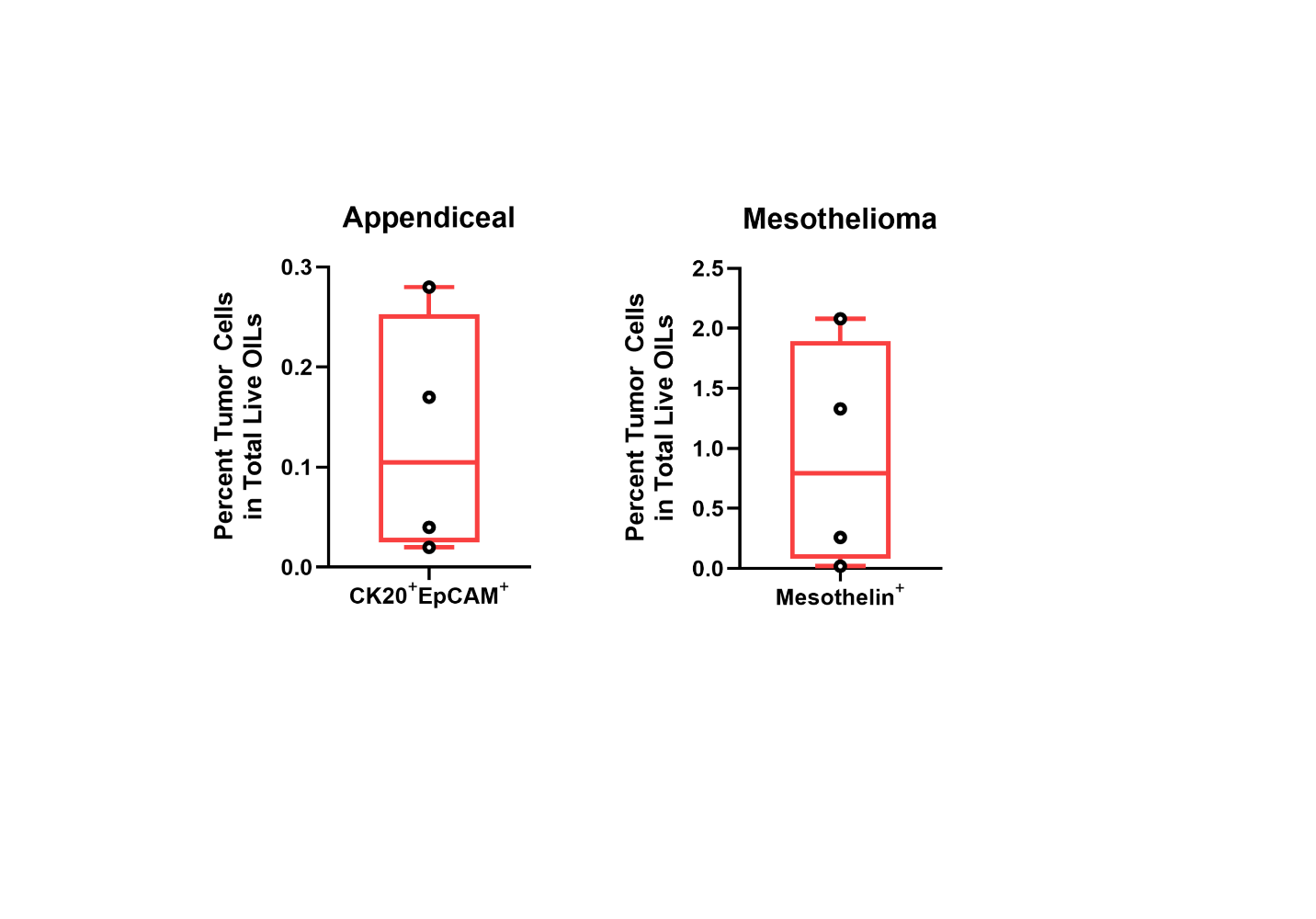

**Fig. S8. OILs show low residual tumor cells at the end of expansion.** Flow cytometry analysis of the percent of total live OILs expressing CK20+EpCAM+ (for appendiceal, left) or mesothelin+ (mesothelioma, right) to identify the abundance of residual tumor cells. Data points represent individual patient specimens, measured in triplicate. Box plots present minimum, median, and maximum values.

**Table S1.** **Cytotoxicity Antibody Panel for Flow Cytometry Analysis.**

| **Catalog Number** | **Antibody Name** |
| --- | --- |
| 563835 | BV650 Mouse Anti-Human CD69 |
| NP242616A94 | Alexa Fluor™ 594 Mouse Anti-Human Cytokeratin 20 |
| Ab196235 | PE Rabbit Anti-Human Mesothelin |
| 11-9983-42 | FITC Mouse Anti-Human HLA-ABC |
| 563792 | BUV395 Mouse Anti-Human CD45 |
| 557834 | APC-Cy™7 Mouse Anti-Human CD8 |
| 560835 | PerCP-Cy™5.5 Mouse Anti-Human CD3 |

**Table S2. Phenotype Antibody Panel for Flow Cytometry Analysis.**

| **Catalog Number** | **Antibody Name** |
| --- | --- |
| 356120 | Brilliant Violet 510™ Anti-Human CD25 |
| 302334 | Brilliant Violet 605™ Anti-Human CD20 |
| 305118 | APC Anti-Human CD66b |
| 351328 | Brilliant Violet 711™ Anti-Human CD127 (IL-7Rα) |
| 304608 | PE/Cyanine5 Anti-Human CD56 (NCAM) |
| 560835 | PerCP-Cy™5.5 Mouse Anti-Human CD3 |
| 557834 | APC-Cy™7 Mouse Anti-Human CD8 |
| 747960 | BV650 Rat Anti-Human CD366 (TIM-3) |
| 568369 | BUV737 Mouse Anti-Human CD4 |
| 743118 | BV786 Mouse Anti-Human CD122 |
| 560922 | PE-Cy™7 Rat Anti-Human CCR7 (CD197) |
| 556626 | FITC Mouse Anti-Human CD45RA |
| 562299 | PE-CF594 Mouse Anti-Human CD45RO |
| 560743 | Alexa Fluor® 700 Mouse Anti-Human HLA-DR |
| 556641 | PE Mouse Anti-Human CD95 |
| 563792 | BUV395 Mouse Anti-Human CD45 |
| 35-4776-42 | PE-Cyanine5.5 Rat Anti-Human FOXP3 |
